# The AberTrap: an open-source, lightweight, programmable sampler of airborne biological particles

**DOI:** 10.64898/2026.09.02.746715

**Authors:** Ruby R. Bye, Andrew P. Detheridge, Mark Neal, Liam George, Hoda M. El-Gharabawy, Bhuvnesh Shrivastava, Philip A. Jennings, Marta Žižek, Ian Izett, Dave Comont, Peter Todd, Shelley Rundle, Tristan Colaço, Matt Wainhouse, Gareth W. Griffith

## Abstract

**Context:** Capturing and identifying airborne biological matter (bioaerosols) enables detailed characterisation of ecological communities, but the cost and weight of existing commercially available active air samplers limits the deployment of multiple devices needed to realise this potential.

**Method:** We present the AberTrap, a lightweight (∼50g), open-source (CC-BY-SA), programmable rotating-arm bioaerosol sampler that can be self-built from common laboratory components and makerspace equipment at low cost (ca £20 per unit, exc. power supply). To characterise the device’s performance, we first explored the influence of rotational speed and paddle design on aerosol capture under controlled conditions, using fluorescein and a range of biological aerosols (4.5-33 µm). To demonstrate its utility for ecological applications, we then validated field performance in a species-rich broadleaf woodland (England, UK), deploying eight samplers over five days and identifying captured fungal aerosols by ITS2 metabarcoding.

**Key results:** Modifying paddle design allowed us to influence particle selection and capture efficiency. Slotted paddles captured significantly more aerosols than plain paddles despite sampling less air, which we traced to the concentration of aerosols at the paddle leading edges. This effect was most pronounced for the smallest particles and outweighs the influence of rotational speed. In the field, 20 daily samples (four samplers over five days) detected 2,431 fungal taxa, 78.9% of the Chao2-estimated total fungal richness. A single sampler captured only 59% of this richness. Continuous 5-day sampling recovered 78% of the taxa detected by daily sampling, indicating only modest paddle saturation over this period. Our time segregated design allowed us to infer dispersal dynamics; 35% of taxa were recorded only on one day, and 19% recorded across all five days. Overall, replicate AberTraps provided high sampling completeness.

**Implications:** The AberTrap provides a low-cost, lightweight and programmable alternative to commercial rotating-arm samplers, enabling levels of spatial and temporal replication impractical with existing devices. Low cost and portability facilitate deployment in remote or resource-limited contexts. Its open-source, adaptable design allows the device to be tailored for applications in biodiversity assessment, pathogen surveillance and ecological monitoring.

## 1. Introduction

### 1.1. The ecological importance of bioaerosols

Airborne travel represents a critical life-history phase for many organisms; permitting reproduction, dispersal and access to fresh resources. Consequently, the air contains a diverse range of biologically derived particles. The best studied of these are microbial propagules (fungal and bacterial spores), and pollen grains, although other types of biological matter such as leaf fragments, animal detritus, lichen soredia and viral particles also become airborne. When suspended in the air, these biogenic particles are usually referred to collectively as bioaerosols (Bohmann & Lynggaard, 2023).

To date, most bioaerosol research has focused on surveillance of disease causing bioaerosols, such as SARS-CoV-2 containing airborne vesicles (Chia et al., 2020; Huang et al., 2024; Kim et al., 2018) or plant pollen, since the pollen of some plant species can trigger rhinitis in susceptible individuals (Brennan et al., 2019; Greiner et al., 2011; Idrose et al., 2020; Kim et al., 2018; Maya-Manzano et al., 2021). Even in non-urbanised settings, when the contents of the air are studied it is most common to trace pathogens, for example *Phytophthora ramorum* through Californian tanoak (*Notholithocarpus densiflorus*) forests (Filipe et al., 2012) or the amphibian-lethal chytrid *Batrachochytrium dendrobatidis* dispersal in fog (Prado et al., 2023). However, bioaerosols are involved in a greater diversity of roles beyond disease causation.

Bioaerosols are essential for ecosystem functioning at multiple scales. They are important for global carbon and water cycling: by acting as condensation nuclei for clouds and translocating carbon across large distances (Bahram & Netherway, 2022). At the local scale, saprotrophic fungi, dispersed primarily by airborne spores, drive nutrient cycling when connected with fresh substrates. Bacteria and fungi influence plant and animal fitness through both mutualistic and pathogenic interactions (Aguilar-Trigueros et al., 2025). Pollen is, of course, an essential part of plant reproduction. Since aerosol release is often transient and environmentally responsive, temporal patterns in bioaerosol composition can reveal organism responses to changing conditions, informing management strategies for both conservation and pathogen surveillance. Similarly, spatial patterns in propagule detection can distinguish whether species distributions reflect dispersal or establishment limitations. These ecological insights require methods able to characterise entire airborne communities.

Environmental DNA (eDNA) approaches allow the simultaneous identification of all taxa present in complex biological assemblages by determining the organismal origin of nucleic acids extracted in bulk from environmental samples (Tedersoo et al., 2022). Metabarcoding or metagenomic approaches are now commonly applied to eDNA isolated from soil and water. However, the application of these technologies to the air lags behind other substrates. Airborne eDNA is typically present at extremely low density, therefore realising the capacity to identify multiple organisms simultaneously from air samples fundamentally depends on the initial capture and concentration of airborne material.

Bioaerosol sampling devices can be broadly categorised as either passive or active. Although cheaper and simpler, passive samplers rely on airflow or rain to deliver aerosols, and are generally limited to qualitative monitoring (Falacy et al., 2007; Grosdidier et al., 2018; Jackson & Bayliss, 2011; Schweigkofler et al., 2004). Active samplers intersect a controlled volume of air, allowing estimation of particle concentrations (Manibusan & Mainelis, 2022). This makes them better suited to quantitative studies, such as pathogen surveillance, public health studies and ecological assessments (Berelson et al., 2025; Ferguson et al., 2019). The most commonly deployed active samplers are either suction or rotation types. Suction types draw the air to the capture medium where particles are separated by either aerodynamic inertia (cyclone and cascade samplers) or pore entrapment (filter samplers), while for rotation type samplers the capture surface is swept through the air, collecting particles by impaction onto the capture surface (Ferguson et al., 2019; Manibusan & Mainelis, 2022; West & Kimber, 2015).

Recent applications of bioaerosol eDNA have demonstrated its capacity to reveal ecological patterns at multiple scales. At the global scale, a network of 47 cyclone samplers deployed worldwide revealed latitudinal gradients in fungal diversity (Ovaskainen et al., 2024). Regional studies using smaller sampler networks have characterised temporal dynamics in airborne pollen across the UK (Brennan et al., 2019) and fungal pathogen distributions in Swedish forest nurseries (Larsson et al., 2025). Several studies have coupled intensive single-location sampling with metagenomics, employing one to three high-intensity devices to characterise urban and agricultural air biomes (Giolai et al., 2024; Gusareva et al., 2020; Qin et al., 2020). In particular, ecological monitoring could benefit considerably from the application of aerial eDNA, by generation of species lists for biodiversity assessments, and tracking community change over time (Giolai et al., 2024; Lynggaard et al., 2022; Polling et al., 2024). Recent proof-of-concept studies have detected terrestrial vertebrates and plants from air samples (Clare et al., 2021; Tournayre et al., 2025) suggesting suitability for landscape-scale biodiversity analysis.

Commercial rotating-arm samplers (Rotorod, Agri Technologies) cost £1,250 and weigh several kilograms, while high-intensity sampling devices (such as the Coriolis micro, Bertin Instruments) cost more than £7,000 (See Suppdata 1.1 for a comparison of the most commonly used active air samplers). However, realising the potential for ecological insight requires both spatial replication across sites and temporal replication within sites, yet the cost and weight of commercially available active samplers restricts the feasibility for concurrently deploying multiple devices.

### 1.2. Aims of this study

Here we present a description and parameterisation of the AberTrap, an open-source, lightweight, programmable sampler of airborne biological particles which can be built at low cost from standard molecular biology lab components using common makerspace equipment. A detailed description of the AberTrap device, including instructions for its assembly and operation can be found in the GitHub repository (https://github.com/RubyBye/AberTrap_air_sampler). To evaluate AberTrap performance for ecological applications, we addressed two aims; **first** to characterise how the operational parameters of rotational speed and paddle design influence capture of diverse bioaerosols under controlled conditions; and **second** to validate its suitability for ecological applications in field tests.

## 2. Materials and Methods

### 2.1. Description and assembly

The AberTrap is an active, rotating arm sampler. Aerosol capture relies on entrapment of propagules in a layer of grease coating the surface of the “paddle”: small rectangular sections of acrylic plastic (29 mm x 7 mm x 2mm; (Fig. 1B), laser-cut from A4-sized sheets. The paddles push-fit into slots on either end of the “arms”: removable, rotating axes printed from flexible TPU filament using a desktop 3D printer (Fig. 1C). The arms push-fit onto the shaft of a small, 2 V solar motor (TruMotion, China), rotation of which is controlled by a Trinket M0 microcontroller (Adafruit Industries, USA) housed within a modified 50 ml screwcap centrifuge (Falcon) tube (Fig. 1A,D). The duration and speed of rotation are programmed by uploading an Arduino sketch via a microUSB port on the microcontroller. The device requires a 5-6V power supply, with typical current draw of 160 mA, which can be provided by 6V lead acid batteries, alkaline battery cells (4×1.5 V AA), or 5V Lithium-ion rechargeable powerbanks. For further details of components and assembly, see (https://github.com/RubyBye/AberTrap_air_sampler).

**Fig. 1.**
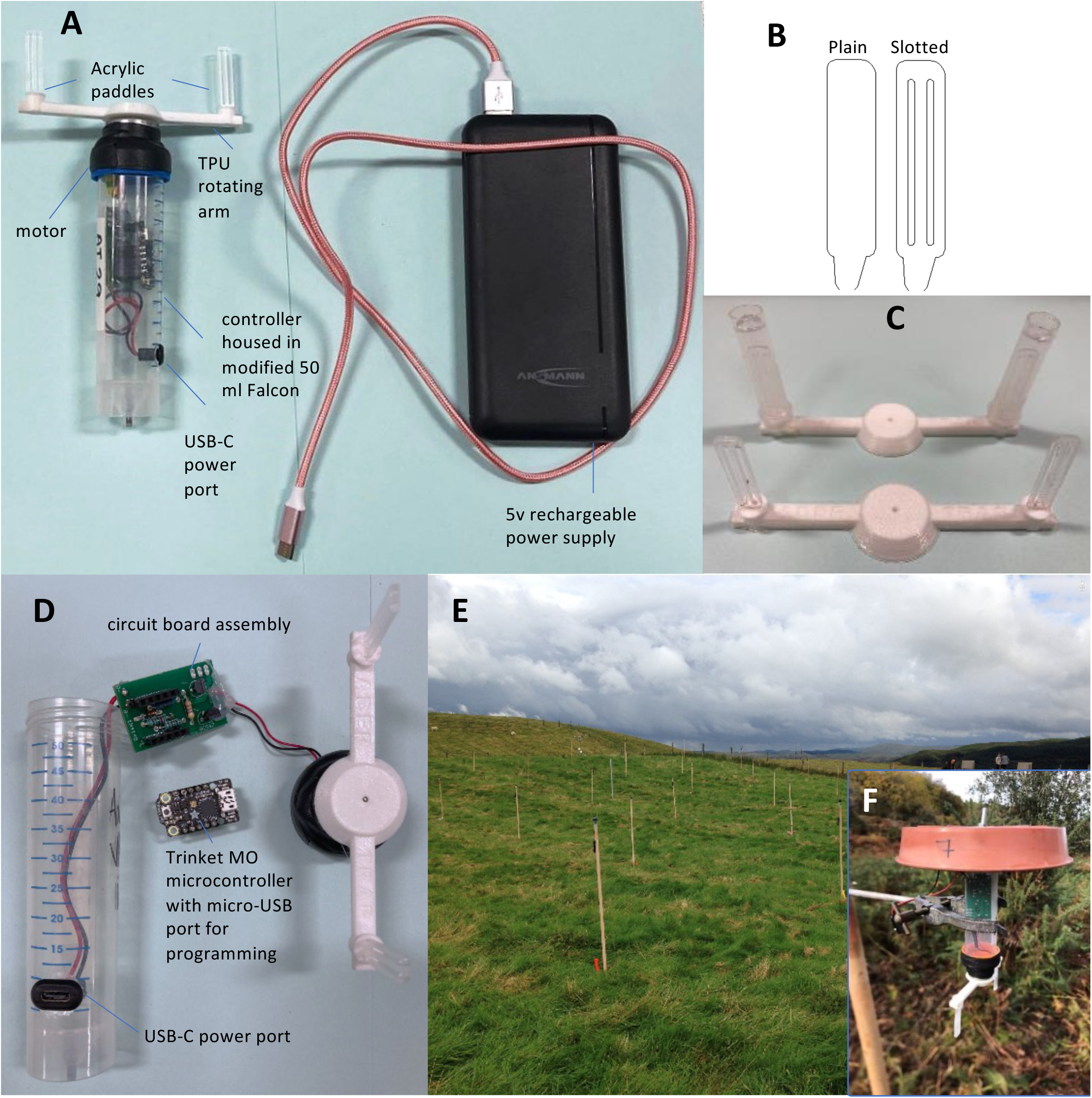
Device description. (**A**) The AberTrap is a rotating-arm type aerosol sampler, comprising (**B**) acrylic paddles mounted in (**C**) a rotating 3D-printed TPU arm, rotation of which is controlled by (**D)** a Trinket microcontroller housed in a 50ml screwcap tube. During sampling, aerosols become embedded in a layer of petroleum jelly on the paddle surface (**B**), where they can be analysed using either microscopy or DNA based methods. (**E**) Low cost of the AberTrap permits affordable use of replicate samplers protected from rain by (**F**) a plastic rain shelter.

#### Paddle preparation

Paddles require coating with grease in preparation of aerosol capture. If using the AberTrap for eDNA analysis then contamination control is of utmost importance: all paddle preparation steps were undertaken in an externally-extracting sterile safety cabinet, in a dedicated clean room in which no amplified nucleic acids are handled. Following laser-cutting, paddles were released from acrylic sheets and washed in sterile, soapy water, to remove any handling residues. Paddles were then rinsed in sterile, deionised water and left to dry in the sterile airflow. Once dry, the paddles were exposed to UV-C light (254 nm ca. 30mJ/cm^2^) for 15 min on a reflective surface to denature any remaining contaminants. Paddles were greased by dipping into a petroleum jelly (Vaseline)-hexane solution (50 mg/mL) using sterile forceps. Dipped paddles were placed on a wire rack to allow hexane evaporation, leaving a thin film of Vaseline (ca. 100 μm thick) covering the entire paddle surface (ca. 100 paddles can be greased in 60 min). The paddles were inserted into the slots on the arm using sterile tweezers. A protective shroud (sterile 2 ml screwcap microcentrifuge tube without lid) was fitted over each paddle (Fig. 1C), securely locked into position on the arm. This shroud prevents exposure of the paddle to the air until its removal on location at the start of sampling. After sampling, shrouds are re-fitted for return to the lab, protecting the paddle from contamination during transit.

### 2.2. AberTrap characterisation

To characterise AberTrap performance, we first determined operational parameters of runtime and sampling rate. We then tested the influence of two main operational elements on aerosol capture: rotational speed and paddle design. Two types of aerosol were used sequentially: 1) non-biological fluorescein aerosols and 2) a mixture of plant and fungal bioaerosols of known sizes.

**Draindown** testing was undertaken with three different power supplies to explore the duration of effective operation. AberTraps were programmed to rotate at 1600 rpm indefinitely using either four AA batteries, a 6V 12Ah lead acid battery or a 5V 20Ah lithium power bank. Voltage was logged until battery depletion below the Trinket cut-off value of 3V.

**The volume of air sampled** (L/min) by the AberTrap was calculated according to the following equation: Swept volume = (paddle area in mm^2^ x (pi x arm diameter in mm) x rotational speed in [rpm])/1×10^6^.

**Paddle designs** compared “plain” and “slotted” paddles (Fig. 1B). For the latter, two parallel slots (each 2mm wide) were made in the paddle face.

**Aerosols** were generated using an airbrush spray gun (Badger Air-Brush Co, Belwood, USA) into a test chamber (0.93 m^3^; Coy Laboratory Products Ltd., USA), with four small desk fans (Kingavon USB Minifan; Model MDUF-113; Shenzhen, China), placed in each corner of the chamber to maintain air circulation. To ensure that no aerosols were collected prior to full dispersal, protective shrouds were removed after a 90 sec delay, and sampler rotation commenced after a 120 sec delay. Capture was assessed over a 10 min runtime. Shrouds were stored in a lidded box during sampling, to prevent accidental aerosol collection. Two types of aerosol were used, as follows.

**Non biological fluorescein aerosol tests** were conducted by aerosolising a fluorescein solution (0.5 mg/ml fluorescein [sodium salt; F6377, Sigma, UK] in 25% glycerol) into the test chamber, as described above. Plain and slotted greased paddles, were tested at two rotational speeds (800 and 1600 rpm). Each AberTrap was fitted with one slotted and one plain paddle. AberTraps were mounted at approx. 60 cm height on retort stands inside a sealed test chamber. Six samplers (= 12 paddles) were spaced 15 cm apart in the centre of the test chamber. The test was repeated three times.

After the 10 min runtime, aerosol distribution on the paddle surface was revealed with UV backlight. Aerosol capture was also quantified fluorometrically. Paddles bearing fluorescein aerosols were placed in 2 ml microcentrifuge tube. To each paddle was added 1 ml Tris (10 mM, pH 8). The tube was vortexed briefly and 200 µl was transferred to a well of a black, flat-bottomed 96 well plate. Each sample was run in triplicate against a set of standards (conc. range: 0.033 - 2.1 nM fluorescein) in a fluorescence plate reader (Hidex Sense, Finland; excitation 485 nm / emission 535 nm; three flashes from above, medium intensity setting).

**Bioaerosol tests** were undertaken to assess the effect of paddle design and rotational speed on the capture efficiency of particles of different sizes. A mixture of four different biological particles (spores and pollen; 4.5-33 µm diam.; Suppdata 2) in roughly equal concentrations were used. To prevent clumping, suspensions were prepared in a non-ionic detergent solution (Nonidet P40; 0.01% v/v). Particle concentrations were determined using a haemocytometer (three replicates) and following mixing, the final suspension contained 5×10^5^/ml of each of the four particle types. The mixed particle suspension (5 ml) was aerosolised (as described above) into the test chamber, delivering a total of ca. 2.5×10^6^ particles of each type.

#### Suppdata 2. Biological particles used for lab parameterising

After a 10 min runtime, particles were quantified *in situ* on the paddle surface with aid of a microscope. To permit the visualisation of particles trapped within the Vaseline, the greased surface of the paddles was coated with clear nail varnish (nitrocellulose/ethyl acetate) diluted 1:1 with acetone. After hardening, the paddles were mounted on a glass slide and particles counted with aid of a 0.25 mm^2^ counting square at 200x magnification. Since particles were unevenly distributed across the paddle width, all particles were counted in a 0.5 mm strip across the full paddle width at the top, mid and base of the paddle (Suppdata 3). Distribution was determined for one plain and one slotted paddle at each 1600 and 800 rpm rotational speeds. This allowed calculation of an ‘edge effect’ defined as the proportion of each type of particle at the leading edge relative to the mean number across the full width of the paddle. This edge effect was then used to streamline counting of the full set of test paddles by only counting particles in a subset of 9 squares, at the leading edges of the paddles. The number of each type of particle collected per paddle was then extrapolated by considering the edge effect and full paddle area.

#### Suppdata 3. Counting schematic

### 2.3. Field testing

Having parameterised the AberTrap under controlled conditions, we next deployed them under field conditions in a semi-natural broadleaf woodland in Somerset, England (50.9653, -3.0714), part of the Wild Neroche site (May 2025). In this experiment, samplers were mounted on a wooden pole inserted into the ground, positioning the sampler at approximately 1.5 m height, protected from rain by a plastic shroud (Fig. 1E). Eight AberTraps were deployed at 10 m spacing for five consecutive days (120 hrs). Four AberTraps ran for the full five day period (= four “long” samples) while on the remaining four samplers the paddles were replaced every 24 hr (= 20 “daily” samples). AberTraps were programmed to cycle between 10 min intervals of 1600 rpm and 400 rpm. After aerosol collection, each paddle was enclosed in a microcentrifuge tube, and stored at 4°C, until the end of the field campaign, when they were frozen at -20°C. Bulk DNA of bioaerosols trapped in the Vaseline was extracted using an in-tube DNA extraction protocol (see https://github.com/RubyBye/AberTrap_air_sampler) under clean-room conditions. DNA was eluted into 100 µl TE buffer.

### 2.4. eDNA metabarcoding

The fungal constituents of the sampled bioaerosol assemblages were explored through metabarcoding of the ITS2 region of the nuclear ribosomal RNA operon, using established methods (see (Detheridge & Griffith, 2021) for more details). Briefly, this involved generating a 300-500 bp portion of the ITS2 region, amplified using a mix of six ITS3 forward primers in equimolar concentrations (Tedersoo et al., 2014), and ITS4 reverse primer, to capture the full range of fungal diversity. TruSeq dual indices were added to sample amplicons, which were then pooled in equimolar concentration, and cleaned, prior to sequencing on an Illumina MiSeq at Wales Gene Park, to generate 2×300 bp paired end reads.

Returned sequences were paired using PEAR (Zhang et al., 2014), then trimmed of primer sequences and quality checked using a bespoke Python script. Sequences passing QC were clustered into unique clusters (Actual Sequence Variants; ASVs) using the UNoise3 algorithm (Edgar, 2016). Taxonomy was assigned to ASVs using the naïve Bayesian classifier (Wang et al., 2007) against a local database built from the latest general eukaryote release of UNITE (v10 2025) (Abarenkov et al., 2010). Relative abundance data was visualised as an excel spreadsheet. Analysis of airborne fungal assemblages was carried out in R Studio v.4.5.1 (R_Core_Team, 2021), using the *vegan* (Oksanen et al., 2016) and *diplyr* (Wickham et al., 2026) packages. Visualisations were made using *ggplot2* (Wickham, 2016).

## 3. Results

To characterise AberTrap performance, we first tested key design features under controlled conditions to evaluate their influence on particle capture, prior to field deployment to assess capture efficiency of fungal bioaerosols.

### 3.1. AberTrap characterisation

The final assembled device (complete with arms and paddles) is approximately 180 mm × 100 mm, weighs ca. 50 g (excluding power supply) and has a component cost of ca. £20 to build (excluding power supply). With a current draw of ca.160 mA at maximum speed of 1600 rpm powering from a 20Ah nominal rechargeable lithium powerbank (ca. 0.4kg, £35) predicts a continuous runtime of approximately 92 hrs (3.8 days; Fig. 2B). This prediction was validated by the draindown test in which the AberTrap was operated at 1600 rpm continuously to battery depletion, with an observed runtime of 95 hrs (Fig. 2A). At slower speed (800 rpm), continuous runtimes extend to 233 hrs (9.7 days; Fig. 2B). Note that voltage to the AberTrap remained stable over this duration, due to the step-up voltage regulator which is a constituent part of the powerpack. Since rotational speed is related to voltage (Suppdata 4), rotational speed remained constant. This contrasts to lead acid (ca. 7 kg for 20Ah 12Ah battery, £30) or AA battery (4x disposable batteries 95g, £3) power supplies, for which voltage dropped continuously (Fig. 2A), which would result in a gradual slowing of rotation. Since lead acid batteries are heavy and their disposal is environmentally problematic, rechargeable lithium power banks with suitable capacity are the preferred power supply for runtimes greater than 24hrs.

**Fig. 2.**
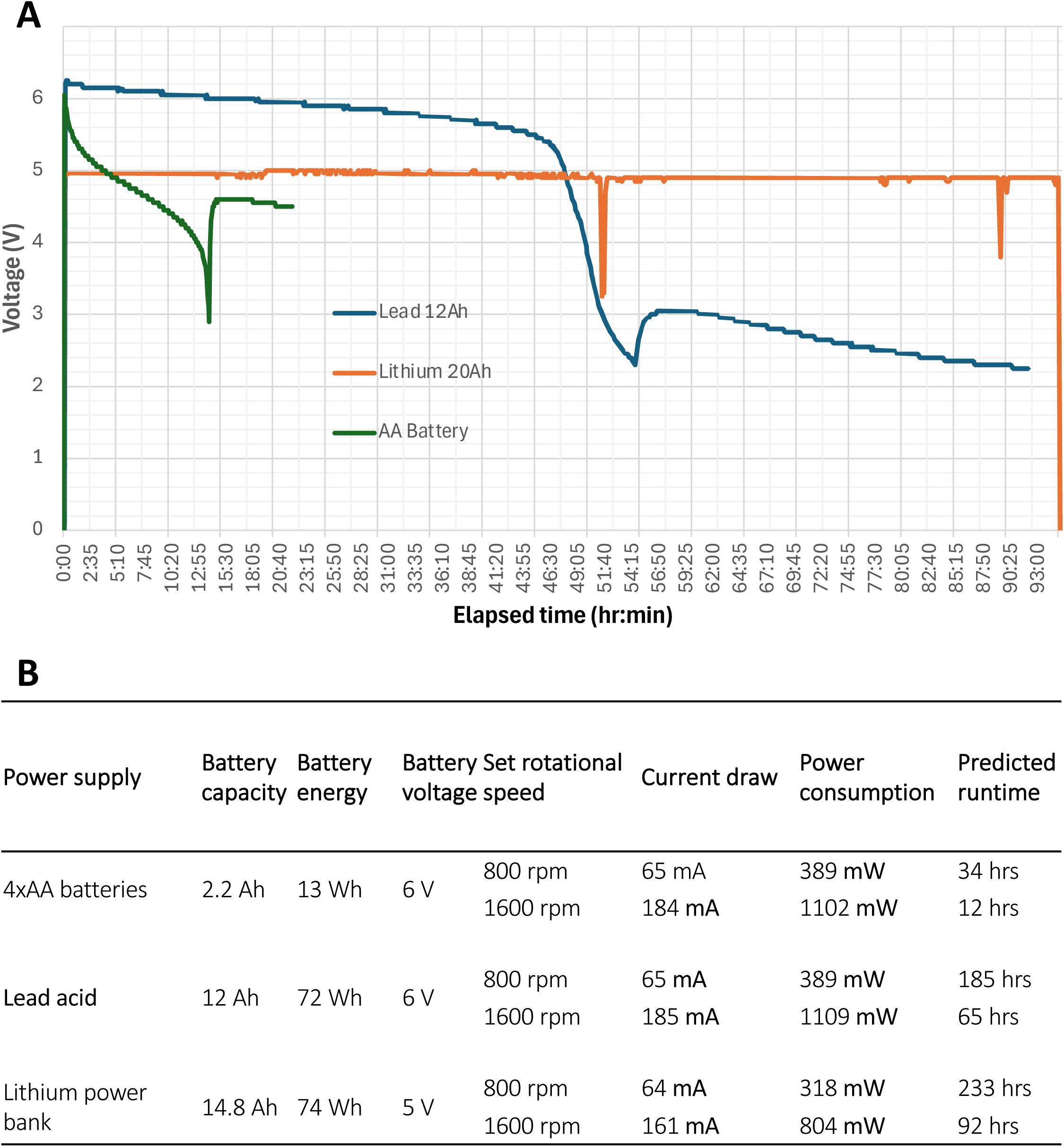
The AberTrap runs for >90hrs on a single 20Ah power bank charge. **(A**) Draindown test showing AberTrap runtime under different power supplies. The sampler was programmed to rotate at 1600 rpm continuously, stopping when the power supply depleted below 3Volts. Note that both 4xAA battery and lead acid power supplies recover some charge after a pause, but drop again below the 3V cutoff voltage. (B) Predicted AberTrap runtimes for different power supplies and rotational speeds

#### Suppdata 4. Voltage influence on speed

### 3.2. Improved capture efficiency of slotted paddles

Aerosol capture under controlled lab conditions showed that both rotational speed and paddle design significantly influenced aerosol capture. Increasing rotational speed increased fluorescein aerosol capture: fewer aerosols (ca. 28% less) were captured at 800 rpm compared to 1600 rpm (Fig. 3B). Paddle design also significantly influenced aerosol capture, but interestingly this relationship did not correspond to the volume of air sampled. Slotted paddles captured three times more aerosols than plain paddles, despite sampling 20% less air volume (72 vs 91 L/min) (Fig. 3A). This difference is statistically significant (two-way ANOVA: speed F=62.38; paddle F=59.29, both p<0.001) (Fig. 3B), suggesting that paddle geometry, not just surface area, determines capture efficiency.

**Fig. 3.**
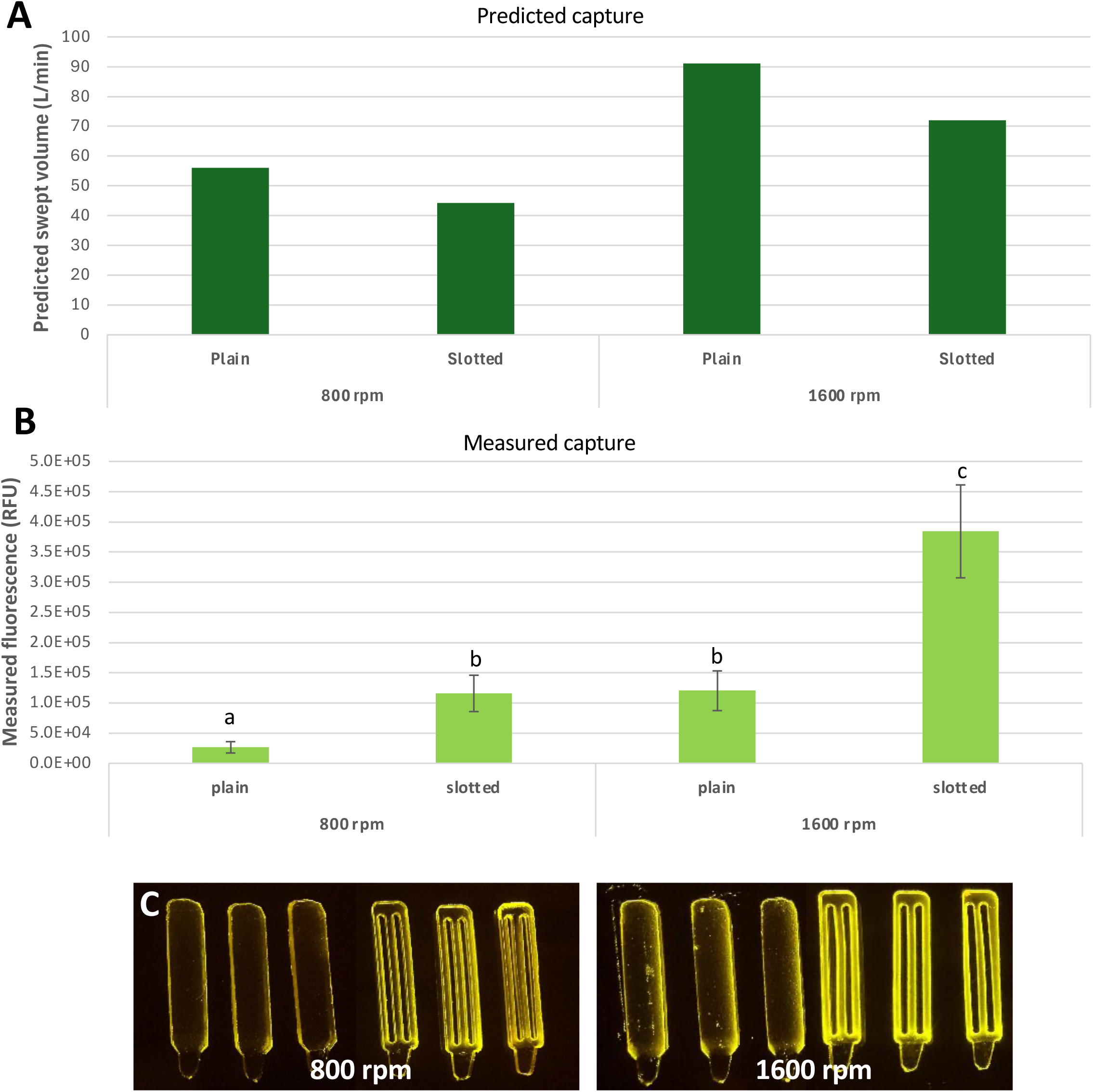
Slotted paddles capture more aerosols than plain, despite smaller swept volume. Three samplers, each bearing one slotted and one plain paddle, were operated simultaneously in a test chamber into which fluorescein aerosols were introduced. **(A)** Predicted swept volume shows that slotted paddles sample a smaller volume of air. **(B)** Despite this, measured fluorescence of the eluates from single paddles showed significantly higher capture by slotted paddles (SE error bars shown, n=9) . (C) The distribution of captured aerosols on the paddle surface reveals a concentration of fluorescence, more apparent at higher speed, at leading edges (curved corner is outermost when rotating).

To explore whether spatial distribution of the particles on the paddle surface could explain why the slotted paddles outperformed plain paddles, we examined fluorescein aerosols in-situ under UV light. This showed fluorescence concentrated at the paddle leading edges of the paddle, dropping within a few mm of the edge (Fig. 3C). Slotted paddles (with three leading edges) therefore had elevated fluorescence across a greater proportion of their width than plain paddles with only a single leading edge. This suggested that the improved capture of the slotted paddles may be explained by the increased number of leading edges. Fluorescein aerosols proved unsuitable to test whether this edge-effect varies with particle size since the liquid aerosol droplets merged on the Vaseline surface to form larger droplets (Suppdata 5) precluding an accurate measurement of their original size.

#### Suppdata 5. Microscope images of fluorescein droplets

### 3.3. Smaller bioaerosols particles concentrated at paddle edges

Using a mixture of bioaerosol particles covering a range of sizes (spores and pollen; 4.5-33 µm diam.; ca. 50-21,000 pg mass range, as calculated via spore volume of sphere and relative density of 1.1; Suppdata 2), we found that the smallest particles (*Calvatia gigantea*, ca. 4.5 µm diam.; *Ganoderma lucidum*, ca. 8 µm diam.) were heavily concentrated at the leading edge(s) (Fig. 4A). Some edge effect could be seen for *Castanea sativa* grains (ca. 17 µm diam.), whilst the largest particles (*Lycopodium clavatum,* ca. 33 µm diam.) were evenly spread across the paddle surface (Fig. 4A). Thus particle size strongly influenced where across the paddle surface the particle became embedded.

**Fig. 4.**
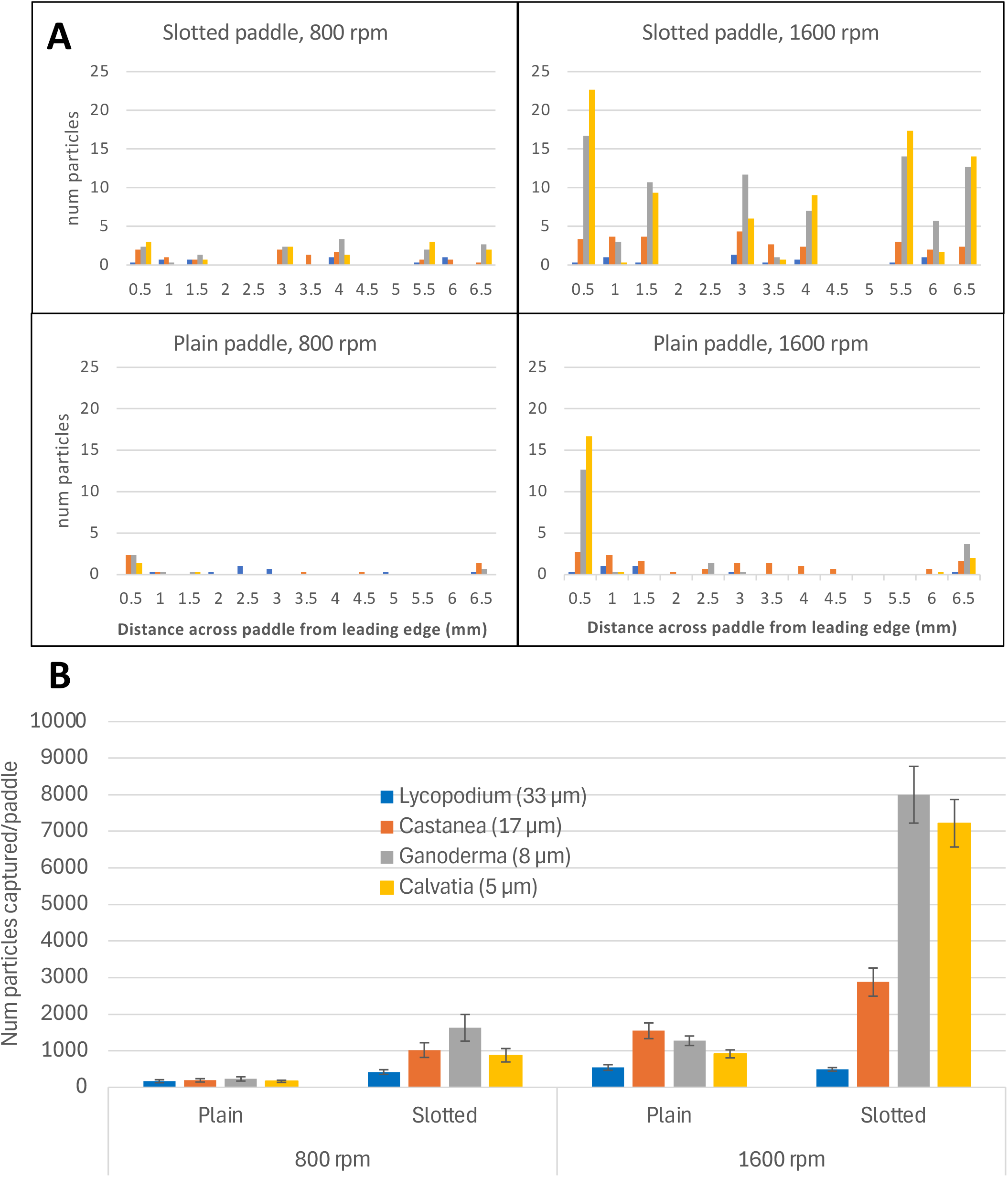
Smaller particles are clustered to the paddle edges. (**A**) Distribution of particles of different sizes across the AberTrap paddle surface. Small particles restricted to paddle edges. The slotted design has more edges and therefore captures greater numbers of smaller particles as compared to plain paddles. (**B**) Increasing rotational speed also improves particle capture. Equal numbers of each particle were introduced to the test chamber. Each test involved three samplers rotating at 800 and 1600 rpm, each bearing one slotted and one plain paddle. Error bars represent SE from three repeats of the test.

This was consistent with the distribution of fluorescein aerosols and indicates an edge-capture phenomenon could explain why slotted paddles showed significantly improved capture of *Castanea*, *Ganoderma,* and *Calvatia* particles at both lower (800 rpm) and faster (1600 rpm) rotational speeds (Fig. 4B). The largest particles (*Lycopodium*) were captured in slightly higher numbers by plain paddles, as might be expected given their fairly uniform distribution across the paddle surface (Fig. 4A) and the greater surface area of plain paddles, although this difference between paddle shapes was statistically significant only at 800 rpm (Suppdata3). Overall, particle size strongly influenced where particles became entrapped on the capture surface. Since smaller aerosols have lower aerodynamic inertia, it is expected that they may divert in the airstream across the paddle and concentrate at the paddle edge. Considered collectively, the results of both fluorescein and particle tests suggest that for a rotating air sampler, paddle shape outweighs rotational speed in influencing capture efficiency.

### 3.4. Field testing of AberTrap

The performance of the AberTrap for exploring natural bioaerosols was assessed by operating several samplers in a semi-natural broadleaf woodland. We used eDNA metabarcoding to identify the bioaerosols captured across a five-day sampling campaign, with four replicate samplers deployed each day (“daily” samples). The weather was stable during sampling, with consistent daily patterns (temp 2-20°C; humidity 48 – 94 %, wind from WNW, 0 – 8.6 km/h gusts, no precipitation).

We found satisfactory sampling completeness with this design. A total of 2,431 unique fungal taxa were identified across all 20 AberTrap “daily” samples (Fig. 5A). A Chao2 estimator (Chao & Lin, 2012), implemented via the *specpool* R function (*vegan* (Oksanen et al., 2016)), was used to predict the total number of taxa that would have been detected by infinite sampling. This analysis predicted a total richness of 3,045 taxa, indicating that 78.9% of the predicted fungal richness present was captured across the five-day period. Parsing taxa according to the day of first detection revealed a declining rate of novel taxon discovery: 1,404 novel taxa were detected on the first day (46% of predicted richness), falling to 138 novel taxa on the fifth day (4.5% of predicted richness; Fig. 5A).

**Fig. 5.**
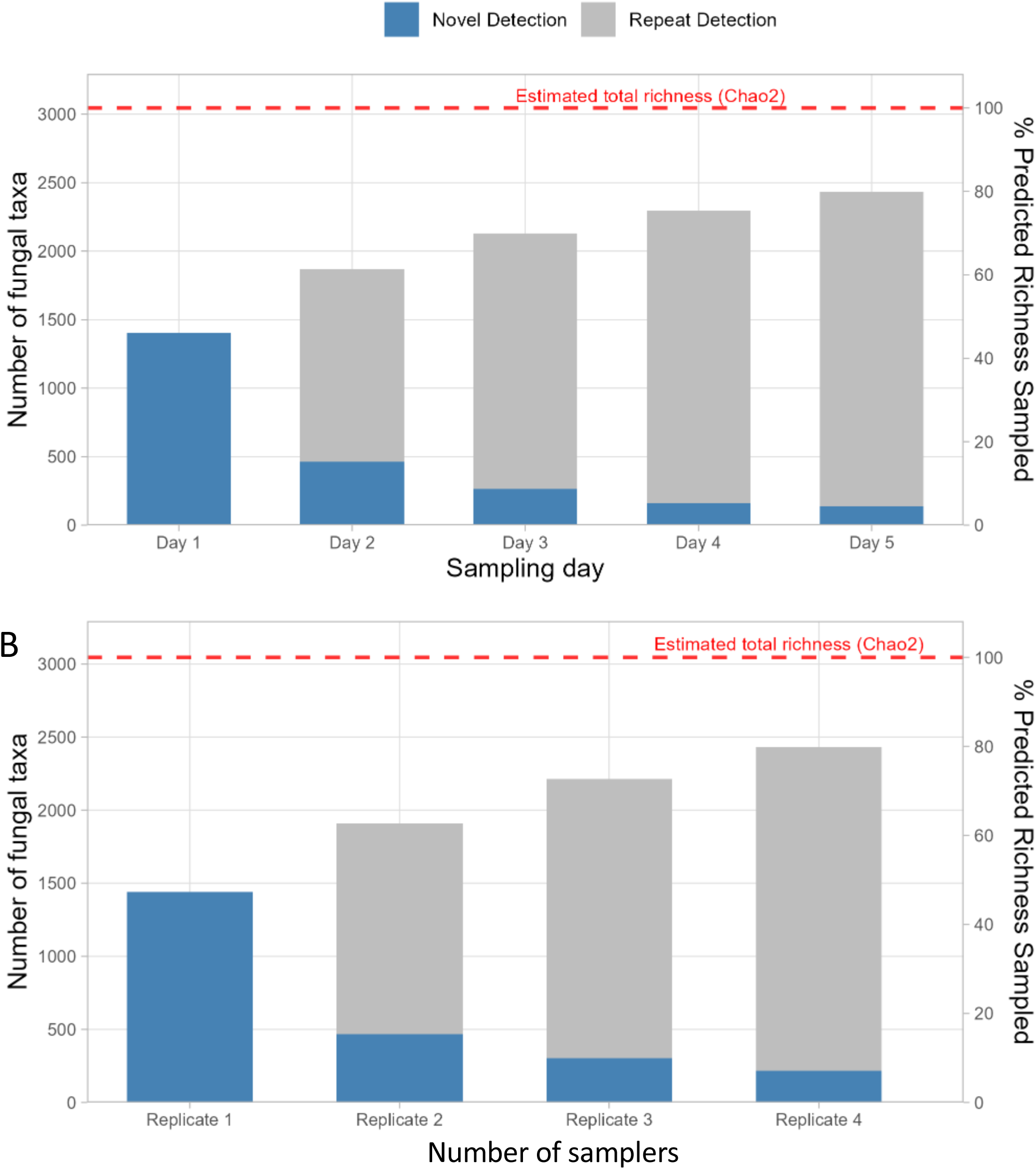
Replicate AberTraps thoroughly sampled the airborne fungal bioaerosols in a semi-natural woodland. Bars show the number of fungal taxa, split into novel and repeat detections (left y axis). Red dashed line indicates the total species richness, estimated with Chao2 from taxon incidence. (**A)** Species accumulation by four AberTraps over five consecutive days (120hrs). Samples were collected at each 24hr interval. Each day adds a declining number of novel species, with cumulative detection at 80% of estimated fungal richness by the fifth day. (**B**) Each replicate sampler, situated at ∼10m spacing, captured additional taxa, with largest gain from adding a second sampler (additional 468 taxa).

We also explored the influence of sampler replication, by parsing taxa according to the sampler on which they were detected (Fig. 5B). Across all five days, a single sampler captured 59% of the detected total number of taxa. Each additional replicate device added additional novel taxa (rep 2: 468 taxa; rep 3: 304 taxa: rep 4: 218 taxa). While this shows a high degree of agreement between the four samplers (placed approx. 10m apart), it highlights the importance of replication. Considered collectively, this indicates that the majority of fungal taxa present were detected by replicated, consecutive daily sampling.

**Detection frequency** (the number of days on which a given taxon was observed) showed a bimodal distribution. The largest group (849 taxa, 35% of total) were observed transiently on only a single day (Fig. 6), while a second peak of 453 taxa (19% of total) were detected repeatedly on all five sampling days. This observation highlights an important technical facet of sampling design; that shorter sampling durations risk missing transient members of the community. More broadly, these detection patterns also permitted categorisation of fungi as either persistent or transient components of the aerial community, providing useful biological information and ecological insight (Fig. 5,6).

**Fig. 6.**
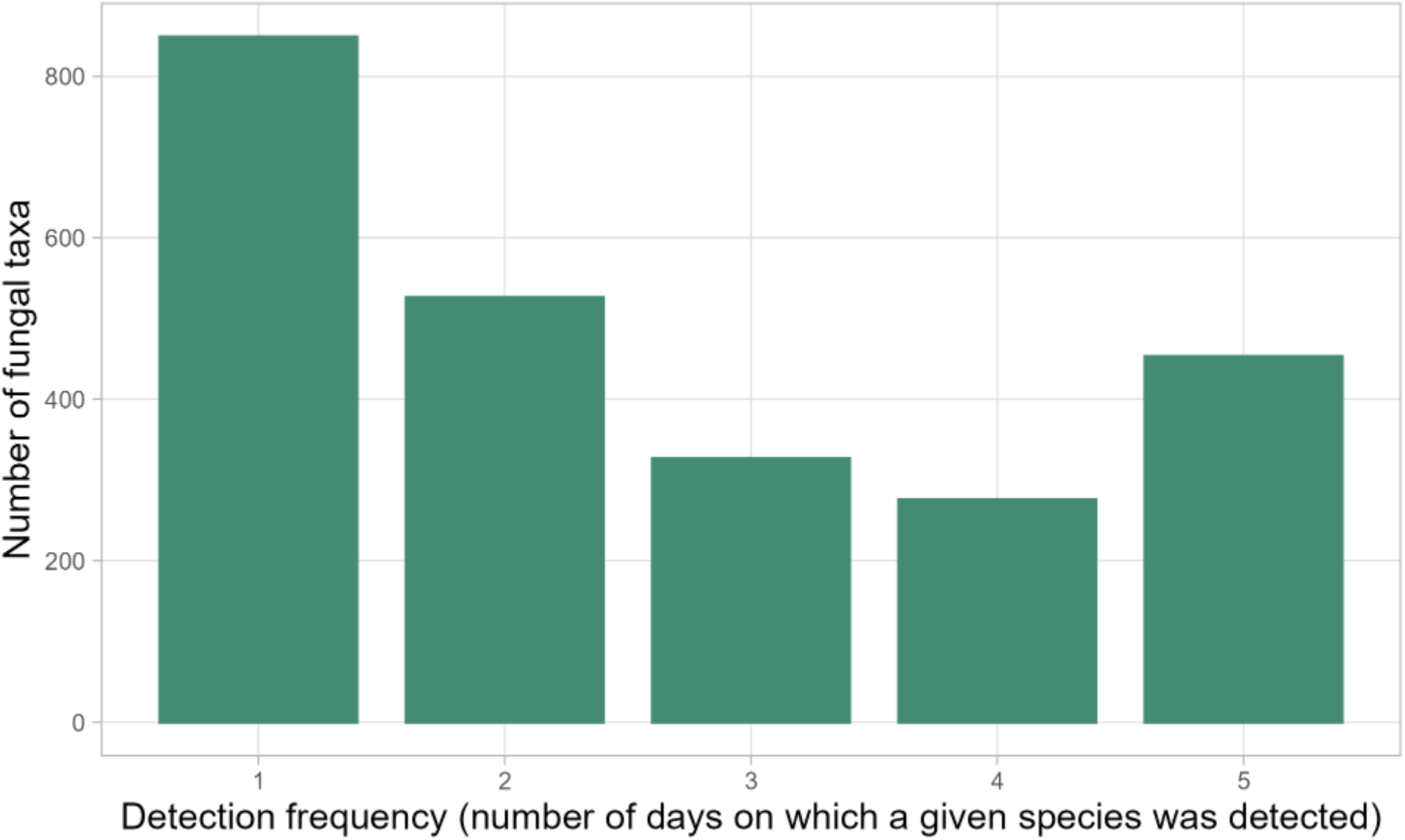
Most fungal taxa were either detected once, or were common. Fungi were categorised according to the number of days on which they were detected. Four replicate air samples were collected each day, for 5 consecutive days Bars show the detection frequency (days) of all detected fungi (n= 2431). The largest group were fungi observed only on a single day (rare fungi; n= 849), with second peak comprising fungi observed consistently across all days (common fungi; n= 453).

Transient species could potentially be more efficiently detected by use of longer sampling durations. However, this approach has upper limits. For example, accumulated material on the paddle surface could reduce stickiness over the longer term, impacting detection at the later stage of particle capture. Alternatively, abundant, common aerosols could dominate those of rarer species, reducing the efficiency of their DNA extraction or sequencing. To empirically test the effect of longer sampling, we operated an additional four samplers continuously for five days (120 hr). These “Long” samples were operated alongside the “Daily” samplers discussed above.

A total of 1,837 taxa were detected in the four “Long” samples. To permit equitable comparison between “Daily” and “Long” samples, all samples were rarified to the same sequencing depth (13,822 sequences for daily samples and 69,108 [5x greater depth] for long samples). After rarefaction, Daily samples yielded 2,014 taxa, and Long samples yielded 1,563 taxa; 78% of the species richness captured by repeated, consecutive daily sampling. Although continuous five-day sampling was slightly less effective at capturing the full spectrum of fungal diversity present, the considerably reduced time spent attending to the samplers may make the reduced number of fungal taxa sufficient, depending on the research objective.

To explore whether detection efficiency declined over time, we identified those fungi present on only a single sampling day (day-exclusive fungi) in the Daily samples. The presence of these fungi acted as timestamps for a given day. Checking for presence of these day-exclusive fungi in the Long samples thus gave a measure of Long sample detection rate, on a day-by-day basis. On average 194 day-exclusive fungi were detected per sampling day (range: 167 to 252 taxa). Long samples detected 42% of the day-exclusive fungi on the first day, falling to 30% of day-exclusive fungi by the fifth day of sampling (Fig. 7), a reduction of less than a third.

**Fig. 7.**
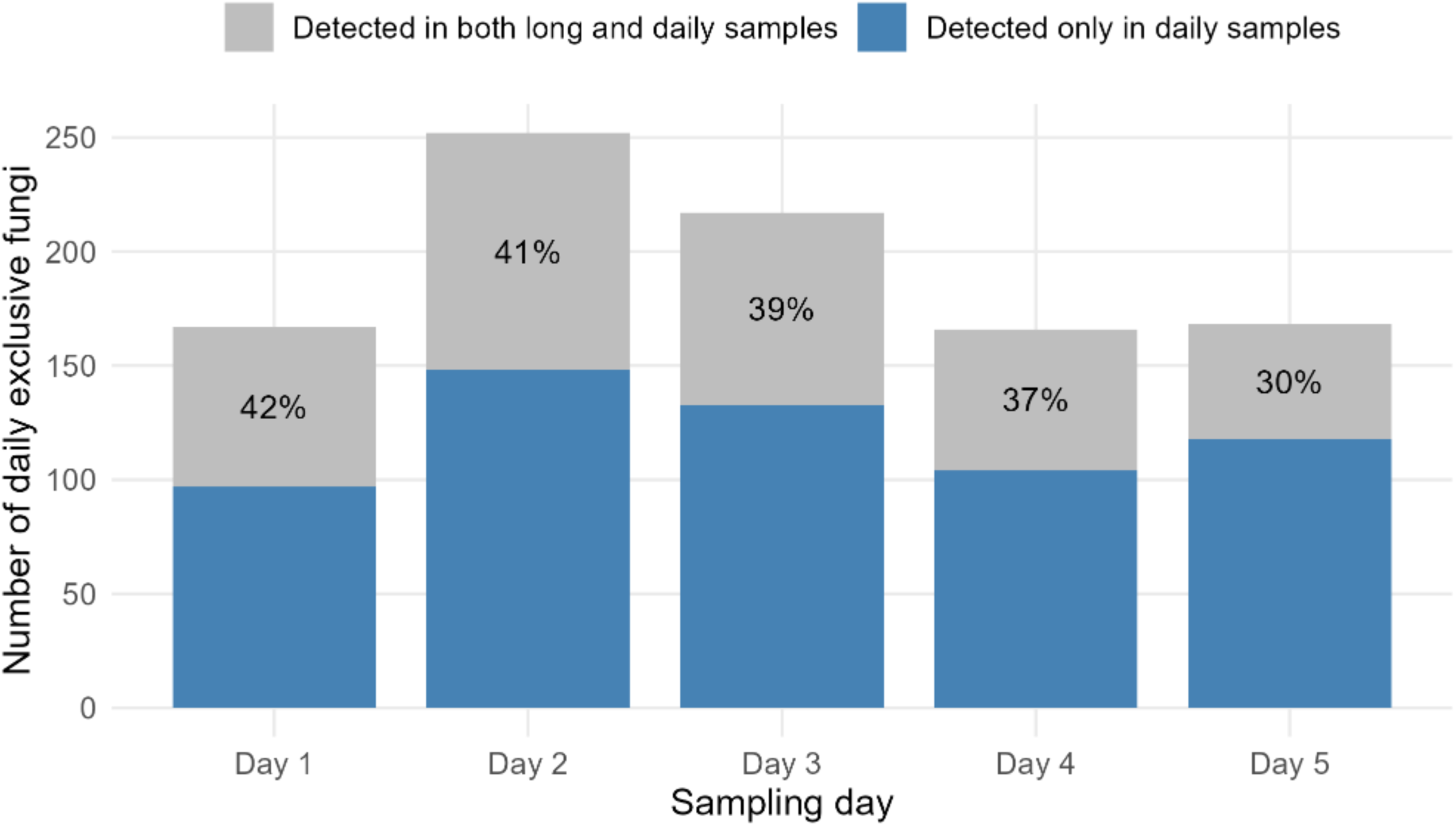
Capture by AberTrap paddles diminished minimally over a five-day runtime. Fungi from daily samples were filtered to those detected on a single day, then matched to the long samples to determine their detection rate on a given day. Bars show the detection of these day exclusive fungi. The proportion of the day exclusive fungi detected by the long samples dropped gradually over the 5 days, reaching 70% of day 1 capture by day 5. Sequence depth was rarified to enable equitable comparison between samples.

## 4. Discussion

Here we present characterisation of the AberTrap, a lightweight rotating-arm aerosol sampler that can be self-built at low cost from common laboratory components using standard makerspace equipment. While other low-cost DIY rotating arm impact sampler designs have been published (Check et al., 2024; Katz et al., 2025) the AberTrap is distinguished by its low power use, programmability, simplicity of operation and open-source design. The design prioritises accessibility through low acquisition cost and portability, enabling deployment of multiple or replicated samplers for ecological studies. Through lab characterisation and field tests, we demonstrate the device’s capacity for comprehensive community sampling, extended operation and detection of ecological patterns previously difficult to investigate with commercial devices.

### 4.1. AberTrap permits sampler replication

The low cost of the AberTrap design makes it feasible to deploy large numbers of traps simultaneously, with its programmable delay permitting synchronisation of operation (Fig. 1E). Remarkably few published aerobiological studies incorporate sampler replication, for example Gygax et al. (2026) due to the high cost of commercially available devices. We found that a single sampler only captured ca. 60% of the taxa detected by four replicates, over a 5-day campaign. Replication increases the depth with which aerial communities can be explored and the reliability of any statistical inferences. While providing similar capture rates to commercial samplers, the AberTrap’s lower cost, weight and size (Suppdata 1) render simultaneous deployment of multiple devices more accessible. When coupled with careful experimental design, multiple samplers can be used to explore spatial and temporal dynamics in the air at finer scales than currently possible with existing devices. Our field tests illustrate how replicated sampling enables ecological insights into community composition, sampling duration, and aerosol temporal dynamics; insights previously restricted to multi-partner collaborative networks (Abrego et al., 2024; Brennan et al., 2019).

### 4.2 AberTraps thoroughly sampled aerial woodland fungi

Operating four samplers simultaneously over five days detected the majority (79%) of predicted fungal richness in a species-rich woodland, demonstrating comprehensive sampling of a complex bioaerosol assemblage. This compares to similar previous studies: 7,065 OTUs (reported as representing “species level”) across 90 x 24hr air samples collected with a Burkard cyclone sampler from urban and forested areas (Abrego et al., 2020) and in a separate study 1,021 distinct taxa were detected in 134 x 24hr Burkard cyclone samples from mostly boreal forest sites (Abrego et al., 2018). The declining rate of novel taxon detection (from 1,404 on day one to 138 on day five) together with high sampling completeness, suggests that most common and abundant taxa present in the area at the time of sampling were detected in our design. More broadly, this demonstrates the capacity of the AberTrap to effectively capture complex bioaerosol assemblages.

While extending the sampling period would likely reveal additional taxa, the returns on sampling effort would diminish unless accompanied by a change in the cohort of aerosols present, triggered for example by meteorological shifts, phenological events or change in host availability. Additionally, practical limits to sampling duration may arise from sample saturation, which we assessed through comparison of one-day versus five-day sampling.

### 4.3. Paddle saturation minimally affects prolonged sampling

Paddle saturation is a recognised limitation of impaction type air samplers (Berelson et al., 2025; Jackson & Bayliss, 2011; West & Kimber, 2015), although empirically determined timeframes of saturation are scarce. Frenz et al.(1999) reported no detriment to sampling for continuous operation up to 72 hours at pollen concentrations below 10 grains/m^3^, although they did not test longer durations. Our five-day continuous sampling test, carried out in spring when pollen and airborne seeds were abundant, showed detection of day-5-exclusive fungi at 71% of those on the first day, a modest decline indicating low paddle overloading. This validates that extended sampling can capture additional taxa without substantial loss to detection capacity over timescales relevant to ecological studies (days to weeks). Whether saturation rates differ across seasons or particle types remains to be determined, though our spring test suggests tolerance even when abundance of non-target aerosols is high.

### 4.4. Time-structured sampling revealed spore dispersal dynamics

Deployment of multiple samplers over multiple days enabled temporal characterisation of the airborne fungal community revealing potentially useful phenological signals. Although airborne fungal communities follow predictable spatial and temporal patterns at the global spatial scale (Abrego et al., 2024), local aerosol mixtures are known to be changeable on the timeframe of hours and days (Fierer et al., 2008; Samake et al., 2019; Womack et al., 2015). In our time-structured design, taxa were categorised as either persistent (detected on all five days; 19% of taxa) or transient (detected on only a single sampling day; 35% of taxa) contributors to the aerial assemblage. This bimodal pattern likely reflects different dispersal sources and strategies: transient detection may originate from intermittent release triggered by environmental change or rare long-distance dispersal. Persistent spores are likely continuous release from local sources.

For biodiversity assessments using aerial eDNA, detection frequency has been suggested as a metric for distinguishing resident species from those passing through. Since most spores deposit locally (Abrego et al., 2024; Norros et al., 2012), repeated detection across multiple days has been proposed to indicate proximity and therefore allows the local community to be identified (Ovaskainen et al., 2020). However, dispersal is also influenced by spore size, with smaller spores (<10 µm) predicted to remain airborne for longer and thus potentially travel further (Norros et al., 2014). The simultaneous detection of hundreds of different spore types through aerial eDNA offers the potential to link detection frequency with spore characteristics for a range of taxa. By keeping detailed records of the weather conditions, AberTraps could therefore be used to explore meteorological factors influencing bioaerosol assemblage composition.

### 4.5. The AberTrap can increase breadth of air eDNA studies

Due to its compact and lightweight design, the AberTrap is well-suited to deployment in distant and/or hard-to-reach field locations. The simplicity of its operation also makes it easily useable by non-expert users. Together, these features offer the possibility of deployment by citizen scientists and/or poorly-resourced scientists in low and middle income countries. We demonstrated this with two successful sampling campaigns undertaken in Egypt and India (Suppdata 6). Four complete AberTraps including sterile, pre-greased paddles were packed into a small plastic box (5×12×17cm) weighing ca. 400 g and posted overseas to the sampling location. AA alkaline batteries were purchased locally. After sample collection, the whole box was returned to Aberystwyth for sample processing and sequencing, demonstrating a straightforward approach to sampling in a broad and distributed range of locations.

#### Suppdata 6. Deployment of AberTrap in distant, inaccessible or challenging locations

### 4.6. Flexible design enables optimisation for research applications

The open-source, modifiable design allows optimisation for specific research contexts through both physical and programmatic modifications. Our tests showed that paddle geometry influences size-selective capture. Paddle shape exerted greater influence on aerosol capture than rotational speed, with edge-to-area ratio determining capture efficiency of different sized particles. The closest commercial comparator, the Rotorod (Suppdata 1) is reported to collect particles only greater than 10 µM (Di-Giovanni, 1998), with simple rod-shaped paddles. We found that paddles with greater edge proportions captured smaller aerosols (5-8 μm) more effectively. This finding has practical relevance for sampling natural bioaerosol communities, in which fungal spores typically span 2-50 μm (Golan & Pringle, 2017) and total bioaerosol size ranges from nanometers to 100 μm (Després et al., 2012). Since rotational speed is also known to influence aerosol capture (Check et al., 2024; Noll & Pilat, 1970), the combination of adjustable paddle shape and programmable speed offers potential for tuning capture characteristics to target particular aerosol size ranges. However, capture efficiency across the full bioaerosol size spectrum remains to be systematically characterised.

Beyond paddle modifications, the rotational program is also user-defined. Extended sampling seems important for comprehensive community sampling; we identified over a hundred novel fungi on the fifth sampling day (Fig. 5A). However, we found a discrepancy between Long sample overall success (78% of daily capture) and the ability to detect the day-exclusive fungi (ca 35% of daily capture) which potentially reveals a risk of omitting rare fungi by prolonged continuous sampling. Day-exclusive fungi, by definition, are episodic components of the aerial community, often present only at low abundance (data not shown). Such rare aerosols may be disproportionately affected by stochastic events during sample processing; they can be lost during nucleic acid extraction or during sequencing library prep, although this is speculative and the principle underlying this observation requires further investigation.

The programmable operation of the AberTrap offers a solution: incorporating pauses into the rotational schedule (e.g., 10 min active - 30 min pause) allows collection to be spread across multiple days while accumulating equivalent biomass to single day sampling. This approach combines the operational simplicity of long continuous sampling, while minimising the risk of sample overloading and associated potential non-detection of rare aerosols. Overall, the rotational program can be optimised for each environment and research goal: comprehensive species lists from dynamic environments may require more intensive sampling, while monitoring target species in stable environments permits longer pauses.

We have made the design files, assembly instructions, and control software available as open-source resources, enabling researchers to adapt the design for specific applications. Paddle designs can be modified using standard free CAD software, while the Arduino-based control system allows programming modifications. We anticipate that community engagement with these resources will drive optimisation for diverse research contexts beyond those explored here. We hope that others will improve on our design to optimise for their own research.

## 5. Conclusion

We present an open-source, programmable, lightweight air sampler that can be self-built at low cost from readily available components. Assembly and operation instructions are provided under Creative Commons license (CC-BY-SA) to facilitate adoption and adaptation by the research community. Through laboratory tests and field validation, we demonstrate that the AberTrap performs comparably to commercial samplers, improving access to bioaerosol research previously constrained by equipment cost. The low cost and portability enable simultaneous deployment of multiple devices, facilitating high-resolution investigation of aerosol spatial and temporal dynamics. Field trials in a species-rich woodland demonstrated the device’s capacity to characterise complex airborne fungal communities with sufficient completeness for ecological analysis and revealed temporal dynamics in community composition. The adaptability of the design, through physical and programmatic modification, enables optimisation for diverse applications including biodiversity assessment, ecological monitoring or pathogen surveillance, with a particular suitability for understudied, hard to reach natural environments.

## Supporting information

Suppdata_1-6

## Acknowledgements

We are grateful to Barry Thomas (Aberystwyth University Physics Department workshop) for invaluable support, Dan Batts (Year in industry student), and Wales Gene Park for providing a sequencing service.

## Authors’ contributions

GWG, RRB and MN conceived the study; device design/optimisation was undertaken by GWG, RRB, MN, PJ, APD, LG, MZ, II, DC and PT; lab and fieldwork was undertaken by RRB, PJ, TC, MW, HME, BS. RRB and GWG wrote the manuscript with input during editing from all other authors. All authors reviewed the final manuscript.

