## Supplementary material for "The AberTrap: an open-source, lightweight, programmable sampler of airborne biological particles": Suppdata_1-6

### Suppdata 1. Commercial sampler comparison

Comparison of the AberTrap with three commercially available samplers was undertaken under controlled indoor conditions (Suppdata 1.1).

**Suppdata 1.1.** Comparison of common air sampling devices. Specifications and costs as October 2025.

| Device name (Manufacturer) | Sampler type | Cost, ex-VAT | Size/Weight | Power requirements | Battery operated runtime | Sampling rate |
|---|---|---|---|---|---|---|
| AberTrap | Rotating impact | £50 (DIY assembly) | 18 x 10 x 3 cm; 50 g excl. power supply (400 g) | 5-6V battery | 7 days (20Ah 5V lithium battery) | 72 L/min |
| Rotorod (Agri Samplers Ltd) | Rotating impact | £1,250 | 45 x 32 x 42 cm; 7 kg excl. power supply (6 kg) | 12V battery, or 35W solar, or 230V mains connection | 7 days (20Ah 12V lead acid battery) | 75 L/min |
| Coriolis µ (Bertin Technologies) | Suction, liquid collection | £7,820 | 22 x 33 x 36 cm; 4 kg incl. battery (inbuilt) | Inbuilt NiMH battery (1 hr runtime), or 230V mains connection. | 10 mins runtime (collection liquid evaporates beyond this) | 100-300 L/min |
| Multi-vial Cyclone Sampler (Burkard Manufacturing Co. Ltd) | Suction, dry collection | £4,032 | 125 x 87 x 77 cm; 24 kg incl. power supply (inbuilt) | Additional 12V battery, or 230V mains connection, or Solar panel | 17 hrs | 16.5 L/min |

Two AberTraps bearing slotted paddles (programmed to rotate at 1600 rpm; 72 L/min) were compared to one Coriolis Micro (set to maximum speed: 300 L/min), one Burkard Multivial Cyclone sampler (16.5 L/min sampling rate) and two Rotorods (2400 rpm; 75 L/min). All samplers were operated concurrently in a test chamber (volume 11.3 $m^3$). Air circulation was achieved by running a floor mounted rotating office fan. The experiment was undertaken three times.

As in earlier experiments (Results 2.2), capture was first assessed by generating fluorescein aerosols within the test chamber. The AberTrap captured similar aerosol quantities to the Rotorod (Suppdata1.2), suggesting that the greater paddle edge length of the slotted AberTrap paddles compensated for its lower rotation speed. The three-fold greater capture of fluorescein by the Coriolis Micro is attributed to its three-fold greater sampling rate, but its use is limited to a maximum 10 min runtime (hence use of 10 min sampling period in this experiment). Capture by the Burkard was low, but detectable, as this device is primarily designed for long (7-day) sampling periods.

The same experiment was then repeated using a mixture of fungal and plant bioaerosols. Between 1 and 14% of the released bioaerosols were recovered by the sampling devices (Suppdata1.3), with the Coriolis capturing the most by a factor of ca. three. Accurate counting of aerosols captured by the Burkard was not possible because many of the *Lycopodium* spores were shattered in the capture tubes, and these fragments obscured other bioaerosols.

It is likely that a high proportion of largest (*Lycopodium*) spores had settled (even with fan circulation) since propagules of this size fall at ca. 240 cm/min in still air (Zanatta et al., 2016) whereas smaller propagules settle more 10 times more slowly (e.g. ca. 24 cm/min for *Ganoderma*; McCarney and West, 2007). The reduced efficiency of the AberTrap relative to Rotorod for the smallest (*Ganoderma* and *Calvatia)* aerosols may be attributable to the higher rotational speed of the Rotorod.


REFERENCE (not cited in main bibliography)

Zanatta, F., Patino, J., Lebeau, F., Massinon, M., Hylander, K., de Haan, M., Ballings, P., Degreef, J., and Vanderpoorten, A. (2016). Measuring spore settling velocity for an improved assessment of dispersal rates in mosses. Annals of Botany 118, 197-206.

McCartney, A., and West, J. (2007). Dispersal of fungal spores through the air. Mycology Series 25, 65-81

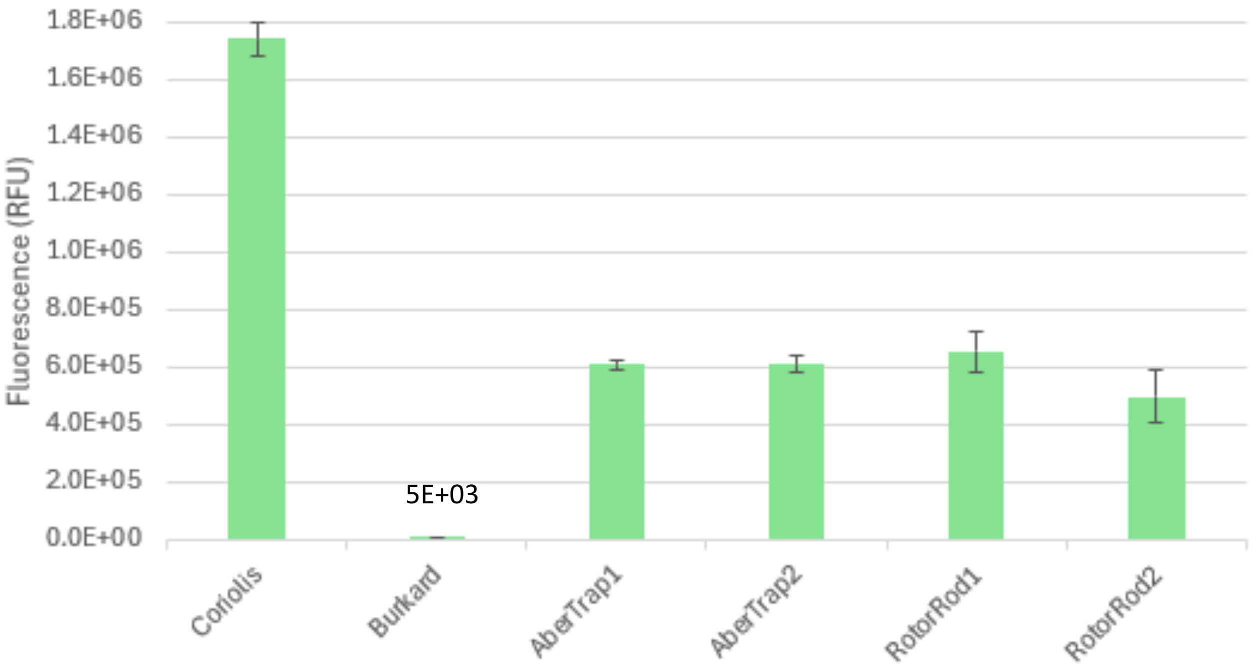


**Suppdata 1.2 Capture of fluorescein aerosols by the AberTrap and commercial samplers.** Barchart shows fluorescence in eluent from each type of sampler. Error bars indicate SE, n=3. Test duration was limited to the Coriolis 10-minute runtime, which likely limited the capture of the Burkard, due to its lower sampling rate.

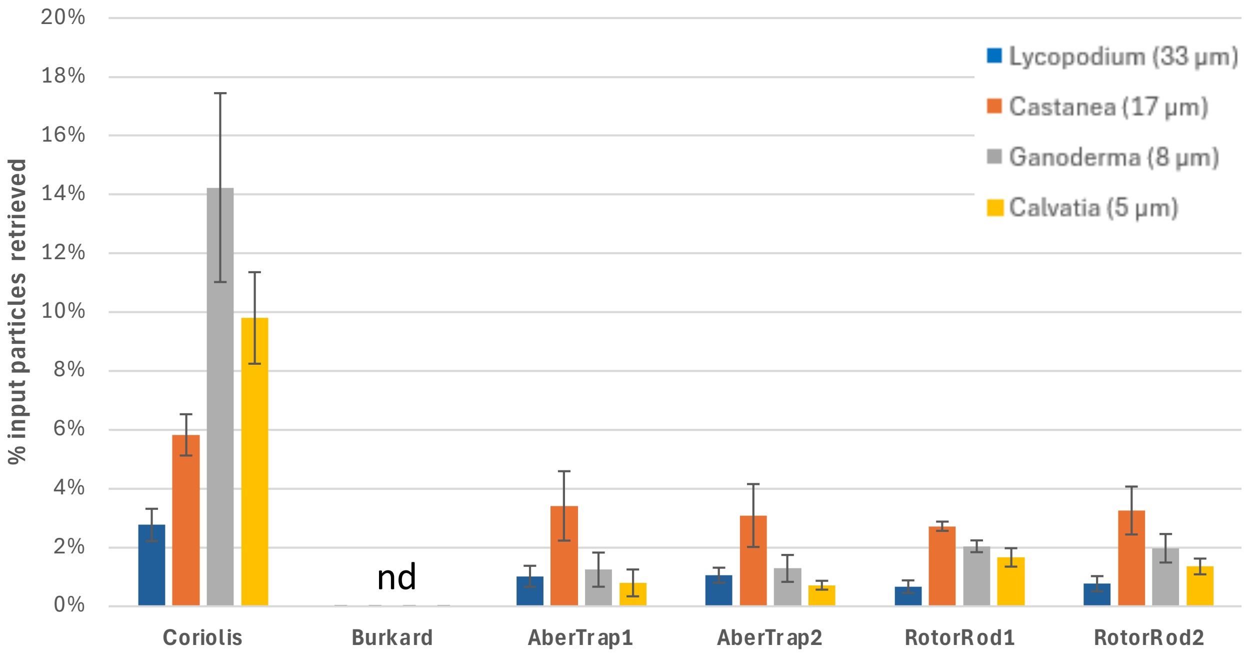


**Suppdata 1.3. Capture of bioaerosols by AberTrap and commercial air samplers.** Barchart shows number of aerosols of different sizes (4.5µm to 33 µm) in 10 min indoor run of AberTrap and three commercial air samplers. Error bars indicate SE, n=3. nd= not detected, not possible to accurately count

#### Suppdata 2. Biological particles used for lab parameterisation

Dimensions and weights for hydrated biological particles used in these tests (measured in water) (Suppdata2.1). Approximate wet weight was calculated using mean propagule diameter and assuming relative density of 1.1g/cm$^3$. Spores/pollen in air will be smaller depending on relative humidity, and when dried will be ca. 10% smaller in diameter (25-35% lower in mass).

**Suppdata 2.1 Dimensions and weights of bioaerosols used for capture tests**

| Propagule | Particle type | Particle description (hydrated) | Approx wet wt.(pg) |
|---|---|---|---|
| *Lycopodium clavatum* | Plant spore (clubmoss) | 28-36 µm, globose/ornamented surface (tetrahedral when dry) | 20,700 |
| *Castanea sativa* | Plant pollen (angiosperm) | 12-15 x 15-20 µm, subglobose (oval when dry) | 1,265 |
| *Ganoderma lucidum* | Fungal spore (bracket fungus) | 9-12 x 6-8 µm, elliptical (finely verrucose) | 420 |
| *Calvatia gigantea* | Fungal spore (giant puffball) | 3.5-5.0 µm, globose (finely verrucose) | 52 |

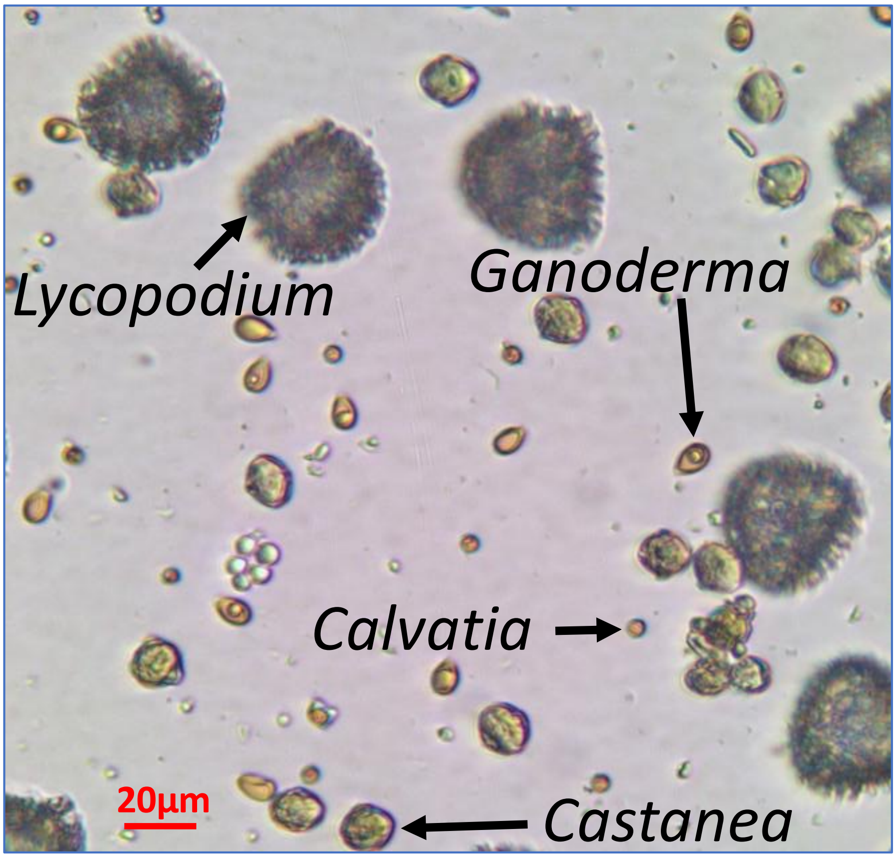

#### Suppdata 3. Counting schematic.

Graphic representation of the schematic in which particles captured on the paddle surface were counted. Particles were unevenly distributed across the paddle width, with greater numbers observed on the leading edges. **(A)** To determine the distribution pattern of each type of particles, numbers of each were counted across the full paddle width. The average proportion at the leading edges was expressed as a proportion of the total number across the full width (Edge Effect factor). **(B)** To streamline counting for test paddles, particles were counted in a subset of 9 squares, at the leading edges of the paddles. The numbers of particles counted here were extrapolated for the whole paddle surface by considering the Edge Effect factor and full paddle area.

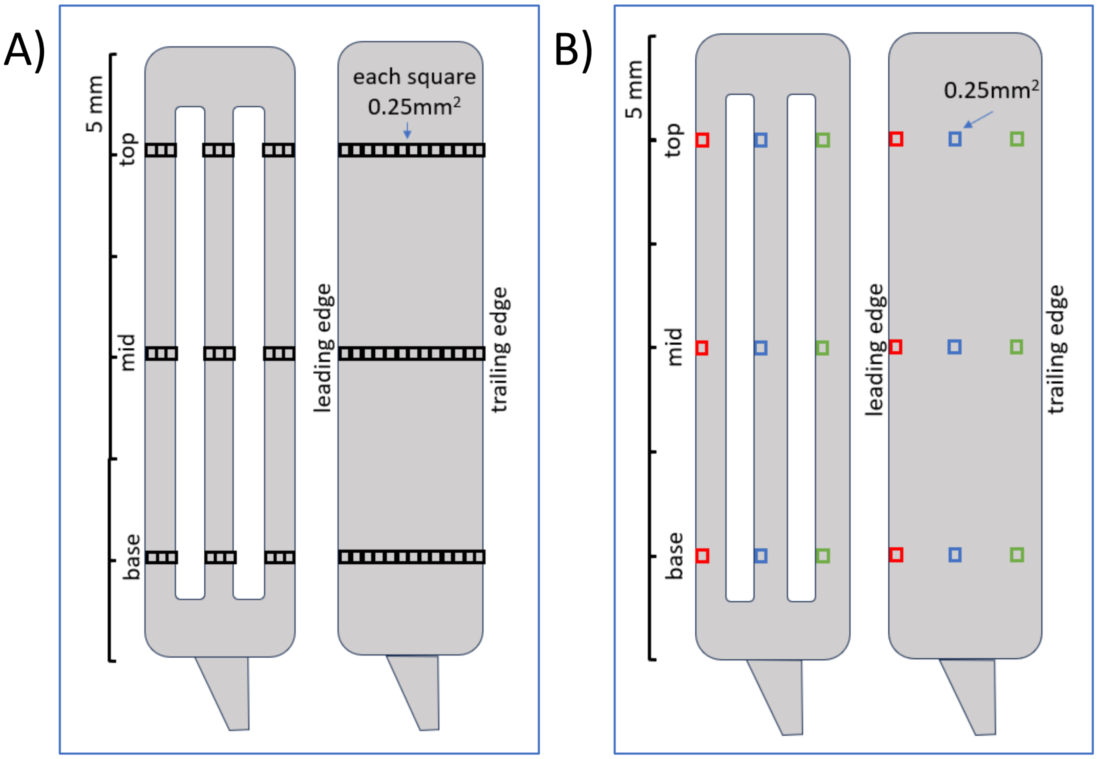


Leading edge

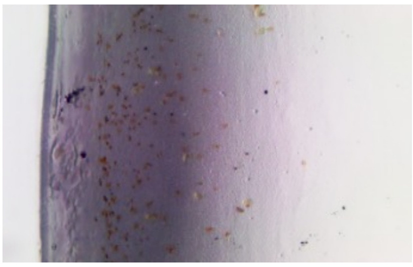

...

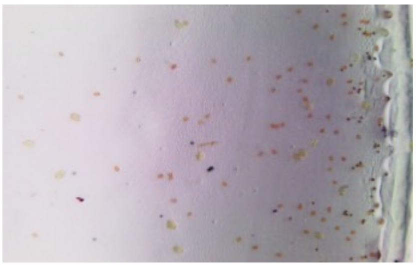

Trailing edge

**Suppdata 3.1.** ANOVA results showing effect of paddle shape ( plain vs slotted)

| | at 800 rpm | | at 1600 rpm | |
|---|---|---|---|---|
| | F | p (same) | F | p (same) |
| *Lycopodium* | 10.41 | 5.27E-03 | 0.3 | 5.82E-01 |
| *Castanea* | 16.31 | 9.52E-04 | 9.1 | 8.06E-03 |
| *Ganoderma* | 14.23 | 1.67E-03 | 73.3 | 2.28E-07 |
| *Calvatia* | 14.7 | 1.46E-03 | 91.7 | 5.00E-08 |

#### Suppdata 4. Relationship between voltage and rotational speed

Rotational speed (RPM, left) is influenced by Voltage, indicating that the declining voltage of the AA and Lead batteries will result in inconsistent speed.

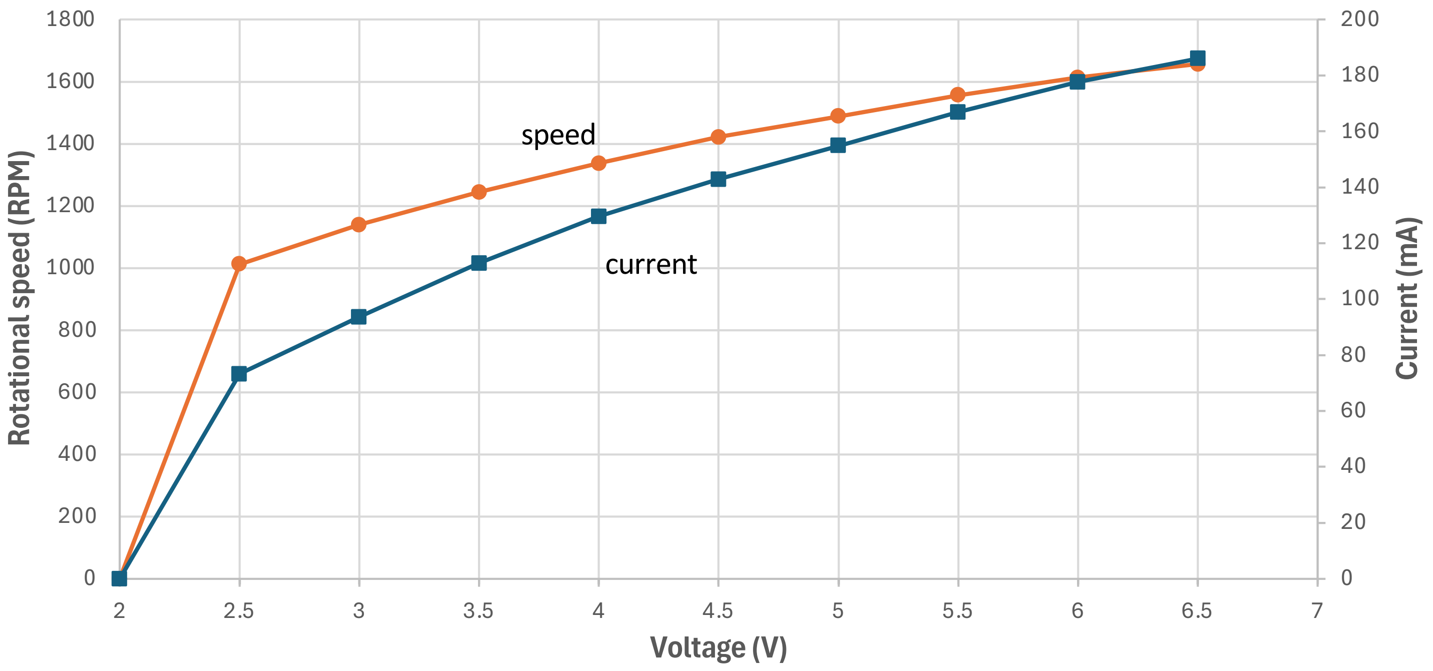

#### Suppdata 5. Fluorescein aerosol positioning

Microscope images showing the layout of fluorescent aerosols captured on the surface of the AberTrap paddles. Note how the aerosol size varies considerably across the surface of the paddle (approx. 3 to 75 µm), with smaller aerosol particles aggregating into larger droplets, most pronounced at the paddle edge (especially leading edge and to a lesser extent the trailing edge). This renders an estimation of airborne aerosol size of fluorescein droplets unreliable using this method.

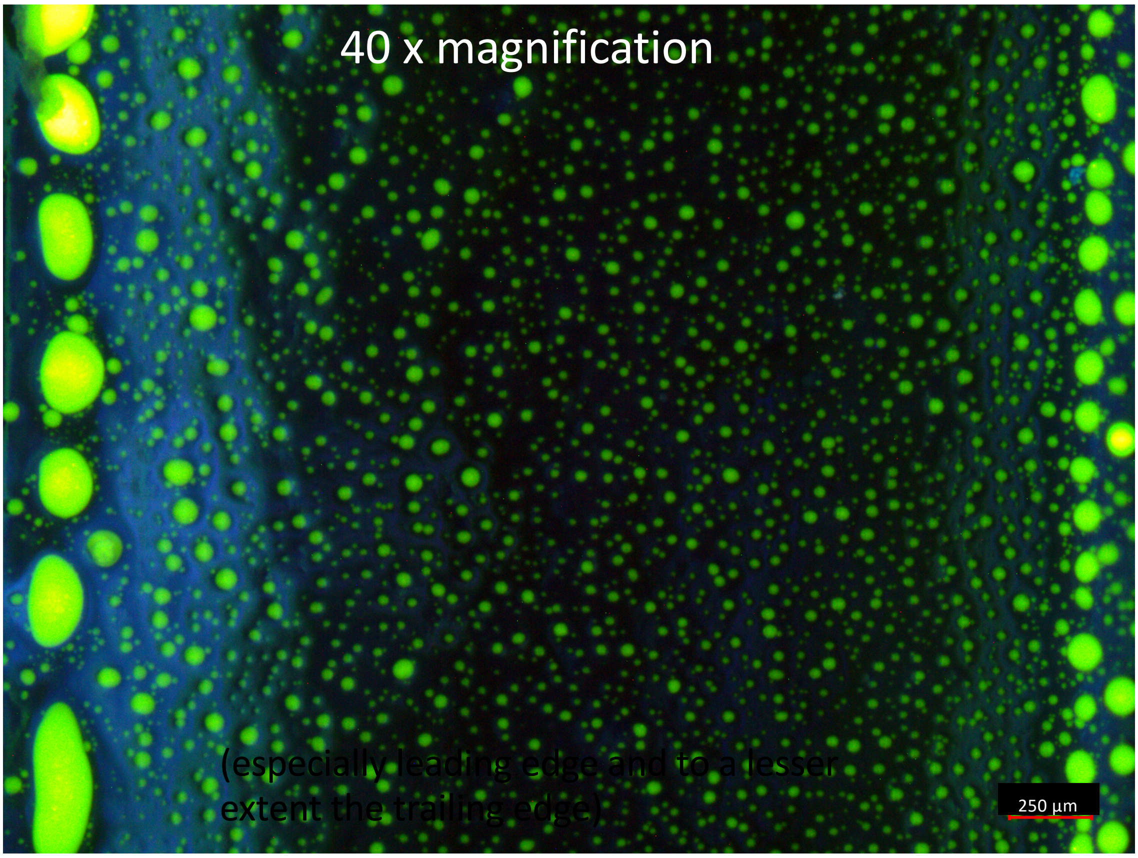


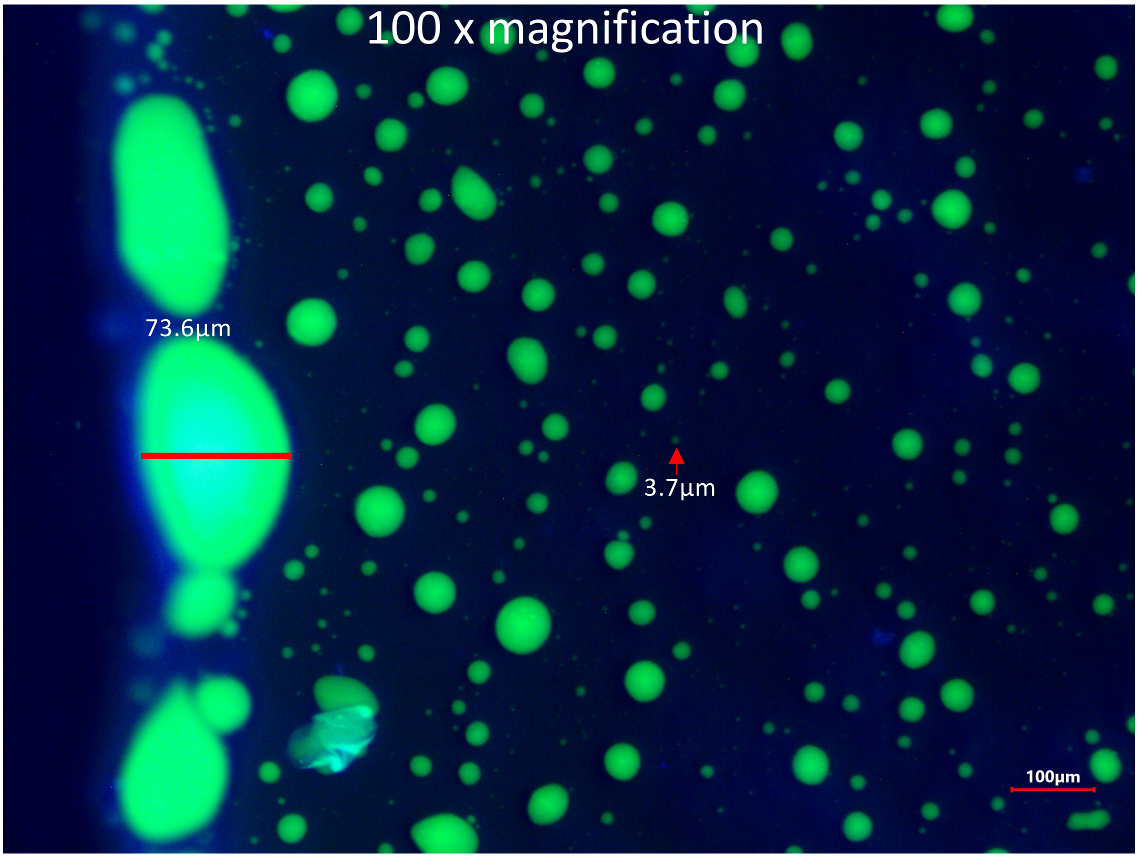

#### Suppdata 6. Deployment of AberTrap in distant, inaccessible or challenging locations

A key feature of the AberTrap is its simplicity of use, compactness and portability. To demonstrate the versatility of the device, we mailed AberTraps to African/Asian collaborators inexperienced in use of the devices. Damietta and New Dehli are at similar latitude but 4300km apart. Sampling both cities was undertaken in mid-winter.

Four AberTraps with associated arms and paddles (pre-greased and stored in 2ml screwcap microfuge tubes) were posted to Dr. Hoda El-Gharbawy in Damietta Egypt (Jan 2015; Suppdata6.1) and Dr. Bhuvnesh Shrivastra in New Delhi, India (Jan/Feb 2016; Suppdata6.2). AA alkaline batteries were purchased locally. AberTraps were deployed in a range of urban, suburban and indoor locations, in sets of four replicates (10 m apart in a square confirguration, started at noon for a 24 hr sampling period (non-slotted paddles, rotating at 800 rpm, noon start/end)(Suppdata6.1). A speed of 800 rpm was used to conserve power from the 4xAA batteries and thus ensure constant speed over 24 hr. Paddles were replaced in their screwcap tubes, stored at 4°C and then mailed (ambient temperature) back to Aberystwyth for analysis.

**Suppdata 6.1.** Details of location and meteorological conditions at the sampling sites

| City | Location | Site code | Lat | Long | Sampling date | Temp range | Wind speed | Wind direction | Paddle numbers | Notes |
|---|---|---|---|---|---|---|---|---|---|---|
| Damietta | University Gardens, grassland | E-UG | 31.4388 | 31.6865 | 26-Jan-15 | 16-19ºC | 9-24 km/h | NE | E01,E03,E05,E08† | †DNA ext from E07 failed but was successful for E08 |
| Damietta | Student residence, rooftop | E-SR | 31.4269 | 31.6567 | 27-Jan-15 | 14-19ºC | 11-24 km/h | NE | E09,E11,E13,E15 | Port Said Airport weather data (50 km away) |
| Damietta | Agricultural, lemon orchard | E-LO | 31.4431 | 31.7796 | 28-Jan-15 | 13-20ºC | 4-24 km/h | W | E17,E19,E21,E23 | https://meteostat.net/en/station/62333 |
| Delhi | Student residence, rooftop | I-SRj | 28.5532 | 77.1744 | 27-Jan-16 | 10-24ºC | 0-11 km/h | N/W | I0I,I03,[I05*],I07 | *Low sequence read number so data discarded |
| Delhi | University Gardens, woodland | I-UGj | 28.5843 | 77.1638 | 28-Jan-16 | 12-24ºC | 0-13 km/h | N/W | I09,I11,I13,I15 | |
| Delhi | Student residence, rooftop | I-SRf | 28.5532 | 77.1744 | 19-Feb-16 | 20-30ºC | 0-13 km/h | W/N | I17,I19,I21,I23** | ** paddle 'thrown' |
| Delhi | Famly home, rooftop | I-FRf | 28.6026 | 77.3179 | 20-Feb-16 | 16-28ºC | 9-20 km/h | W | I25,I27,I29,I31 | New Delhi Safdarjung Airport weather data (8-15 km away) |
| Delhi | Famly home, indoor | I-FIf | 28.6026 | 77.3179 | 21-Feb-16 | 14-26ºC | 9-28 km/h | W | I33,I35,I37,I39 | https://meteostat.net/en/station/42182 |

Only one paddle was lost from the 32 pairs deployed (dislodged from the spinning arm, causing the sampler to become unbalanced and fail to sample air correctly), so 31 intact pairs of exposed paddles were returned. A coating of fine atmospheric dust was apparent on the second paddle of several of the samples sets (Suppdata6.3), notably from one New Delhi rooftop (I-SRj/I-SRf). As noted in the main text the second paddle is kept in reserve both as a backup and for later microscopic analysis of captured propagules etc. until eDNA matabarcoding is completed. For example, initial DNA extraction for paddle E07 failed, but was successful from its pair E08.

Protocols for DNA extraction from paddles follow those described in the main text. eDNA metabarcoding was undertaken using the LSU D1 variable region and sequenced using a IonTorrent PGM, as described in Detheridge et al. (2016/2020). The bioinformatic pipeline is also described in Detheridge et al. (2021).

Across the 31 samples (12 from Egypt and 19 from India, a total of 2,442,402 sequence reads were obtained, of which 230,794 sequences were rejected during quality control and a further 24,355 rejected as non-fungal. The mean number of fungal sequences per sample was 72,666 (range 204,829-20,731). One sample (I05; I-SRj) gave a low number of reads (7,270) so was excluded from subseqent analyses.

Analysis of the fungal communities detected from samples located at three locations in Damietta on successive days were distinct from each other (Permanova Pseudo P=0.002; Suppdata6.1C), with replicate paddles clearly grouping together and 89/1% of the total variancer explained in the two primary PCs. Of the five sample sets obtained in New Delhi, the University Garden and Indoor Family home samples were the most distinct (Permanova Pseudo P=0.0001; SuppFig2B), with the three rootop samples ordinating more similarly. The fungal assemblages captured in the AberTraps deployed on the Student Rooftop three weeks apart (I-SRj/I-SRf) ordinated slightly apart but were not significantly different (Permanova Pseudo P=0.092).

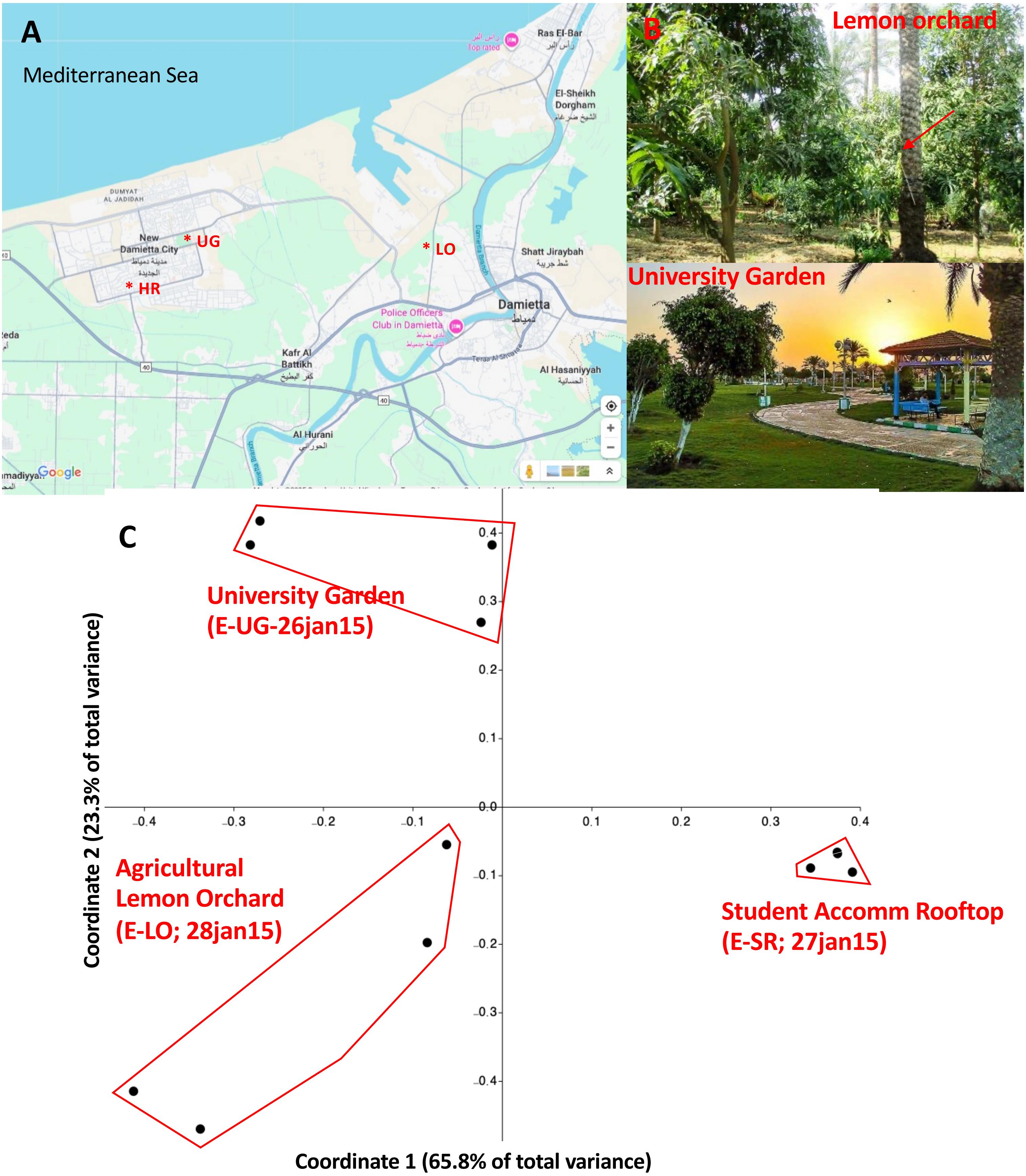


**Suppdata 6.2**: **(A)** AberTrap sampling locations in Damietta, Egypt (Jan 2015). Sampling was undertaken on successive days at three contrasting sites in the city, as detailed in SuppTab1. **(B)** Sampling in lemon orchard, with AberTrap arrowed. Weather was dry all three days. **(C)** Principal Coordinates Analysis (PCoA) ordination of fungal assemblages capture during AberTrap sampling, with percentage of total variance accounted for in first two PC's shown on each axis.

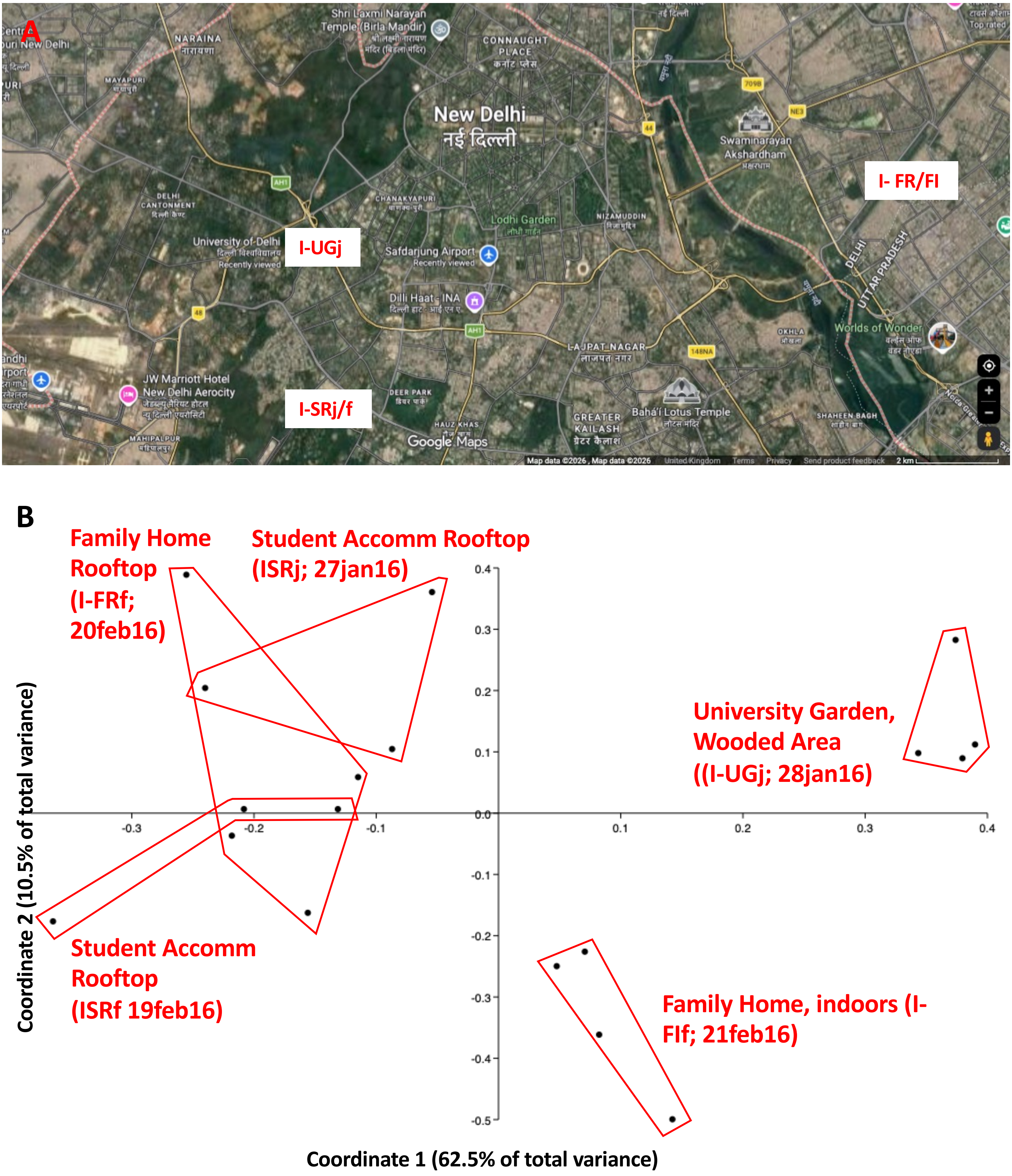


**Suppdata 6.3. (A)** AberTrap sampling locations in New Delhi, India (Jan/Feb 2016). Sampling was undertaken at three contrasting sites in the city, as detailed in SuppTab1, with repeat sampling at two of these sites three weeks apart. **(B)** Principal Coordinates Analysis (PCoA) ordination of fungal assemblages capture during AberTrap sampling, with percentage of total variance accounted for in first two PC's shown on each axis.

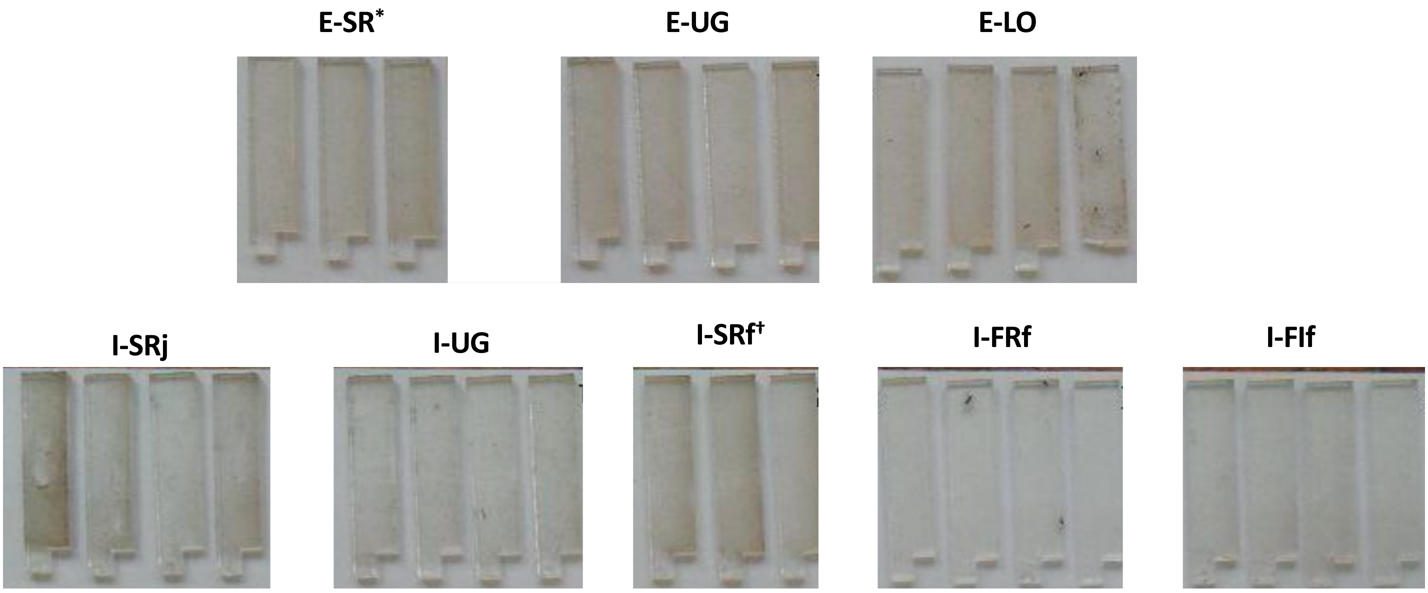


**Suppdata 6.4.** Macroscopic appearance of AberTrap paddles after 24hr duration sampling in Damietta, Egypt (Jan 2015) and Delhi, India (Jan/Feb 2016). Note the accumulation of dust on some sets of samples, especially roof of student residence (I-SRj/f). For the E-SR and I-SRf sample sets, one paddle was dislodged during sampling. (E-SR* -only three paddles shown as reserve paddle was used for repeat DNA extraction; **I-SRf†** -paddle was thrown during sampling).

1136 OTUs were detected amongst the Damietta samples and 726 OTUs in New Delhi samples, of which 470 were common to both cities, including the 24 most abundant OTUs across the whole dataset.

By default, the RDP classifier for the LSU D1 region of the ribosomal RNA locus used here identified species to genus level only and for many genera, this region is insufficiently variable to identify OTU's to species level, for example for several of the fungi most commonly found in air samples more generally (Abrego et al., 2024; e.g. *Alternaria*, *Epicoccum*, *Pleospora*, *Davidiella*, *Cladosporium* spp.; SuppTab6.2 green).

However, for some groups of fungi, species level identification is possible via manual curation (BLAST searching) of retrieved sequeces. For example, *Graphiola phoenecis* (Exobasidiales), a common pathgogen of date palms (genus *Phoenix* ) (Cole 1983) was >1,000-fold more abundant in Damietta than in New Delhi, especially in the lemon (and date palm) orchard (SuppTab6.2; yellow). In New Delhi it was only detected in the indoor household samples (in which palm matting was present).

Similarly spores of the recently described *Peronospora a* (Mu et al., 2024)(OTU41 in SuppTab6.2 orange; mean 2.3% relative abundance), an oomycete pathogen of *Plantago* spp. were abundant from the Damietta rooftop sample but absent from New Delhi samples. The host specificifty of *Peronospora* spp. on *Plantago* spp. is poorly understood (Mu et al., 2024) but it is likely that the host for the propagules detected here was *Plantago ovata* (psyllium/desert Indianwheat), a crop plant widely grown in the Nile Delta. It abundance in only rooftop samples (15 m above ground level) suggests that the source of the propagules was from more distant agricltural areas in the Nile Delta where the crop is widely grown.

In contrast, one unidentified xerophilic *Aspergillus* species (OTU75 in SuppTab6.2 grey; in subgenus Aspergilli; subsection Restricti; Sklenář et al., 2017), was 16th most abundant taxon across New Delhi sampling (mean relative abundance: 0.4%) but absent from Damietta samples. Several other taxa also showed contrasting relative abundance between Damietta and New Delhi samples, including a *Xerochrysium* species (likely *X. coryli*; OTU57 lilac) a xerophytic fungus (Pitt et al., 2013) associated with dried food

**Additional references (not cited in main bibliography)**

Cole, G.T. (1983). *Graphiola phoenicis*: a taxonomic enigma. Mycologia, **75(1)**, pp. 93-116.
Detheridge, A.P., Brand, G., Fychan, R., Crotty, F.V., Sanderson, R., Griffith, G.W., and Marley, C.L. (2016). The legacy effect of cover crops on soil fungal populations in a cereal rotation. Agriculture, Ecosystems & Environment **228**, 49-61.
Detheridge, A.P., Cherrett, S., Clasen, L.A., Medcalf, K., Pike, S., Griffith, G.W., and Scullion, J. (2020). Depauperate soil fungal populations from the St. Helena endemic *Commidendrum robustum* are dominated by Capnodiales. Fungal Ecology **45**, 100911. 10.1016/j.funeco.2020.100911.
Ghoneem, K., Ali, A.A. and El-Baz, S.M. (2007). First record of blond psyllium (*Plantago ovata*) Root rot and wilt diseases in Egypt. Journal of Plant Production, **32(9)**, pp.7273-7287.
Mu, M., Choi, Y.J., Kruse, J., Crouch, J.A., Ploch, S. and Thines, M. (2024). Single host plant species may harbour more than one species of *Peronospora*–a case study on *Peronospora* infecting Plantago. Persoonia-Molecular Phylogeny and Evolution of Fungi, **52(1)**, pp.94-118.
Pitt, J.I., Lantz, H., Vinnere Pettersson, O. and Leong, S.L.L. (2013). *Xerochrysium* gen. nov. and *Bettsia*, genera encompassing xerophilic species of *Chrysosporium*. IMA fungus, **4(2)**, pp.229-241.
Sklenář, F., Jurjević, Ž., Zalar, P., Frisvad, J.C., Visagie, C.M., Kolařík, M., Houbraken, J., Chen, A.J., Yilmaz, N., Seifert, K.A. and Coton, M. (2017). Phylogeny of xerophilic aspergilli (subgenus *Aspergillus*) and taxonomic revision of section Restricti. Studies in Mycology, **88**, pp.161-236.

**Suppdata6.4.** Top 200 taxa (across all samples) detected in the AberTrap sampling from Damietta and New Delhi, ranked by cumulative total relative abundance. Five commonest genera (green) were detected in all samples but some species showed distinctive distributions (yellow/orange/grey/pink, with higher abundance in blue) ,as described in main text.

| Phylum | Class | Order | Family | Genus | Count | Cumul Tot | Mean | Max | Min | E01 |
|---|---|---|---|---|---|---|---|---|---|---|
| Ascomycota | Dothideomycetes | Pleosporales | Pleosporaceae | OTU2 (Alternaria_spp.) | 30 | 803.4% | 26.78% | 44.43% | 2.774% | 43.92% |
| Ascomycota | Dothideomycetes | Pleosporales | Didymellaceae | OTU 1 (Epicoccum spp.) | 30 | 759.4% | 25.31% | 55.68% | 3.314% | 4.36% |
| Ascomycota | Dothideomycetes | Pleosporales | Pleosporaceae | OTU6 (Pleospora spp.) | 30 | 271.4% | 9.05% | 36.34% | 1.482% | 6.733% |
| Ascomycota | Dothideomycetes | Capnodiales | Davidiellaceae | OTU5 (Davidiella spp.) | 30 | 162.2% | 5.41% | 13.01% | 1.179% | 10.75% |
| Ascomycota | Dothideomycetes | Capnodiales | Davidiellaceae | OTU9 (Cladosporium_spp.) | 30 | 90.0% | 3.00% | 7.58% | 1.023% | 1.517% |
| Basidiomycota | Tremellomycetes | Filobasidiales | Filobasidiaceae | OTU10 (Filobasidium_spp.) | 30 | 66.0% | 2.20% | 7.28% | 0.463% | 4.417% |
| Basidiomycota | Exobasidiomycetes | Exobasidiales | Graphiolaceae | Graphiola phoenecis | 15 | 62.58% | 2.09% | 23.81% | 0.000% | 0.007% |
| Ascomycota | Eurotiomycetes | Eurotiales | Trichocomaceae | Penicillium | 30 | 39.06% | 1.30% | 17.88% | 0.015% | 0.195% |
| Ascomycota | Eurotiomycetes | Eurotiales | Trichocomaceae | OTU19 (Aspergillus spp.) | 30 | 33.15% | 1.11% | 3.49% | 0.024% | 1.347% |
| Basidiomycota | Ustilaginomycetes | Ustilaginales | Ustilaginaceae | Ustilago | 30 | 28.06% | 0.94% | 7.98% | 0.051% | 0.457% |
| Ascomycota | Eurotiomycetes | Eurotiales | Trichocomaceae | Eurotium_spp.:OTU20 | 30 | 27.82% | 0.93% | 2.30% | 0.107% | 1.790% |
| Ascomycota | Dothideomycetes | Pleosporales | Phaeosphaeriaceae | OTU 487 (Phaeoseptoriella spp.) | 27 | 24.88% | 0.83% | 2.34% | 0.000% | 0.047% |
| Basidiomycota | Wallemiomycetes | Wallemiales | Wallemiaceae | Wallemia | 30 | 24.78% | 0.83% | 2.71% | 0.052% | 1.362% |
| Basidiomycota | Microbotryomycetes | Sporidiobolales | Sporidiobolaceae | Sporobolomyces;OTU 18 | 30 | 23.98% | 0.80% | 6.15% | 0.031% | 3.718% |
| Pseudofungi | Oomycetes | Peronosporales | Peronosporaceae | Peronospora | 25 | 18.96% | 0.63% | 3.97% | 0.000% | 0.015% |
| Pseudofungi | Oomycetes | Peronosporales | X | OTU 24 | 25 | 16.80% | 0.56% | 4.39% | 0.000% | 0.291% |
| Basidiomycota | Pucciniomycetes | Pucciniales | Pucciniaceae | Puccinia | 30 | 16.32% | 0.54% | 0.99% | 0.161% | 0.648% |
| Ascomycota | Dothideomycetes | Capnodiales | Capnodiales incertae sedi | Devriesia | 17 | 13.36% | 0.45% | 8.48% | 0.000% | 0.171% |
| Fungi incertae sedi: | Mucoromycotina | Mucorales | Rhizopodaceae | Rhizopus | 29 | 12.33% | 0.41% | 3.20% | 0.000% | 0.200% |
| Ascomycota | Dothideomycetes | Pleosporales | Massarinaceae | Massarina | 30 | 10.48% | 0.35% | 1.20% | 0.020% | 0.048% |
| Ascomycota | Eurotiomycetes | Eurotiales | Trichocomaceae | Eupenicillium | 30 | 10.00% | 0.33% | 1.41% | 0.049% | 0.495% |
| Ascomycota | Eurotiomycetes | Eurotiales | Trichocomaceae | OTU 57 (Xerochrysium colyli) | 10 | 9.96% | 0.33% | 2.90% | 0.000% | 2.904% |
| Basidiomycota | Agaricomycetes | Agaricales | Psathyrellaceae | Coprinopsis | 24 | 9.63% | 0.32% | 2.29% | 0.000% | 0.104% |
| Ascomycota | Leotiomycetes | Helotiales | Sclerotiniaceae | Botrytis_ssp:OTU 43 | 30 | 9.57% | 0.32% | 0.78% | 0.011% | 0.171% |
| Pseudofungi | Oomycetes | Peronosporales | Peronosporaceae | **OTU 41 (Peronospora a)** | 6 | 9.45% | 0.32% | 3.78% | 0.000% | 0.000% |
| Fungi incertae sedi: | Mucoromycotina | Mucorales | Mucoraceae | Mucor | 19 | 8.92% | 0.30% | 4.74% | 0.000% | 0.057% |
| Ascomycota | Dothideomycetes | Dothideales | Dothioraceae | Columnosphaeria | 27 | 8.46% | 0.28% | 2.21% | 0.000% | 2.206% |
| Ascomycota | Dothideomycetes | Pleosporales | Pleosporaceae | Cochliobolus | 30 | 7.75% | 0.26% | 0.84% | 0.006% | 0.579% |
| Ascomycota | Dothideomycetes | Dothideales | Dothideales incertae sedi | Selenophoma | 17 | 7.23% | 0.24% | 1.51% | 0.000% | 0.000% |
| Basidiomycota | Tremellomycetes | Filobasidiales | Filobasidiaceae | Naganishia | 30 | 7.14% | 0.24% | 0.89% | 0.032% | 0.102% |
| Ascomycota | Eurotiomycetes | Eurotiales | Trichocomaceae | Aspergillus | 30 | 7.02% | 0.23% | 1.21% | 0.017% | 0.163% |
| Ascomycota | Eurotiomycetes | Eurotiales | Trichocomaceae | OTU 75 (Aspergillus s. Restricti) | 17 | 6.69% | 0.22% | 1.17% | 0.000% | 0.000% |
| Basidiomycota | Cystobasidiomycetes | Erythrobasidiales | Erythrobasidiaceae | OTU 67 | 30 | 6.46% | 0.22% | 0.74% | 0.050% | 0.143% |
| Basidiomycota | Agaricomycetes | Agaricales | X | OTU 4288 | 16 | 6.39% | 0.21% | 3.08% | 0.000% | 0.004% |
| Ascomycota | Dothideomycetes | Capnodiales | Mycosphaerellaceae | OTU 82 | 28 | 6.15% | 0.21% | 4.30% | 0.000% | 0.023% |
| Ascomycota | X | X | X | OTU 263 | 11 | 5.86% | 0.20% | 5.50% | 0.000% | 0.000% |
| Fungi incertae sedi: | Mucoromycotina | Mucorales | Mucoraceae | Amylomyces | 25 | 5.41% | 0.18% | 3.98% | 0.000% | 0.312% |
| Basidiomycota | Tremellomycetes | Cystofilobasidiales | Cystofilobasidiaceae | Cystofilobasidium | 28 | 5.38% | 0.18% | 0.99% | 0.000% | 0.721% |
| Basidiomycota | Microbotryomycetes | Sporidiobolales | Sporidiobolales incertae | Rhodotorula 4 -Sporidiobolales | 30 | 5.24% | 0.17% | 0.44% | 0.040% | 0.220% |
| Ascomycota | Eurotiomycetes | Eurotiales | Trichocomaceae | OTU 62 | 29 | 5.13% | 0.18% | 1.16% | 0.014% | 0.014% |
| Basidiomycota | Agaricostilbomycetes | Agaricostilbales | Agaricostilbaceae | Bensingtonia | 13 | 4.78% | 0.16% | 1.16% | 0.000% | 0.034% |
| Ascomycota | Dothideomycetes | Capnodiales | Mycosphaerellaceae | Ramularia | 11 | 4.65% | 0.15% | 2.05% | 0.000% | 0.008% |
| Ascomycota | Saccharomycetes | Saccharomycetales | Saccharomycetaceae | Kloeckera_lindneri | 29 | 4.58% | 0.15% | 1.25% | 0.000% | 0.006% |
| Ascomycota | Dothideomycetes | Pleosporales | Phaeosphaeriaceae | Phaeosphaeria | 29 | 4.39% | 0.15% | 2.31% | 0.000% | 0.168% |
| Ascomycota | Dothideomycetes | Capnodiales | Mycosphaerellaceae | OTU 124 | 7 | 4.28% | 0.14% | 2.78% | 0.000% | 0.006% |
| Ascomycota | Leotiomycetes | Erysiphales | Erysiphaceae | Erysiphe | 27 | 4.23% | 0.14% | 0.54% | 0.000% | 0.000% |
| Ascomycota | Dothideomycetes | Dothideales | X | OTU 64 | 30 | 4.20% | 0.14% | 0.40% | 0.017% | 0.315% |
| Basidiomycota | Cystobasidiomycetes | X | X | OTU 162 | 13 | 4.09% | 0.14% | 1.24% | 0.000% | 0.028% |
| Basidiomycota | Agaricomycetes | Polyporales | X | OTU 78 | 10 | 4.00% | 0.13% | 3.14% | 0.000% | 0.000% |
| Ascomycota | Dothideomycetes | Pleosporales | X | OTU 785 | 25 | 3.92% | 0.13% | 0.30% | 0.000% | 0.027% |
| Ascomycota | Dothideomycetes | Capnodiales | Mycosphaerellaceae | OTU 128 | 12 | 3.87% | 0.13% | 1.94% | 0.000% | 0.019% |
| Ascomycota | Dothideomycetes | Pleosporales | X | OTU 101 | 29 | 3.85% | 0.13% | 0.40% | 0.000% | 0.184% |
| Ascomycota | Dothideomycetes | Pleosporales | Didymellaceae | OTU 1396 | 30 | 3.74% | 0.12% | 0.88% | 0.009% | 0.014% |
| Ascomycota | Dothideomycetes | Pleosporales | X | OTU 561 | 24 | 3.59% | 0.12% | 0.38% | 0.000% | 0.014% |
| Basidiomycota | Exobasidiomycetes | Malasseziales | Malasseziaceae | Malassezia | 28 | 3.58% | 0.12% | 0.51% | 0.000% | 0.009% |
| Ascomycota | Dothideomycetes | Mycosphaerellales | Teratosphaeriaceae | Hortaea;OTU 47 | 18 | 3.46% | 0.12% | 1.41% | 0.000% | 0.069% |
| Ascomycota | Dothideomycetes | Pleosporales | Didymellaceae | OTU 1446 | 24 | 3.29% | 0.11% | 0.35% | 0.000% | 0.013% |
| Ascomycota | Pezizomycetes | Pezizales | Pyronemataceae | Lasiobolidium | 14 | 3.28% | 0.11% | 2.56% | 0.000% | 0.080% |
| Ascomycota | Leotiomycetes | Helotiales | X | OTU 85 | 19 | 3.26% | 0.11% | 1.05% | 0.000% | 0.130% |
| Ascomycota | Saccharomycetes | Saccharomycetales | Saccharomycetaceae | Wickerhamomyces_anomalus | 22 | 3.25% | 0.11% | 0.70% | 0.000% | 0.582% |
| Ascomycota | Dothideomycetes | Dothideales | X | OTU 115 | 24 | 3.16% | 0.11% | 0.55% | 0.000% | 0.033% |
| Basidiomycota | Tremellomycetes | Trichosporonales | Trichosporonaceae | Trichosporon | 27 | 3.12% | 0.10% | 0.76% | 0.000% | 0.012% |
| Ascomycota | Dothideomycetes | Pleosporales | X | OTU 129 | 29 | 3.07% | 0.10% | 0.40% | 0.000% | 0.007% |
| Ascomycota | Saccharomycetes | Saccharomycetales | Saccharomycetaceae | Geotrichum_silvicola | 30 | 2.92% | 0.10% | 0.43% | 0.013% | 0.120% |
| Ascomycota | Pezizomycetes | Pezizales | Pezizaceae | Iodophanus | 26 | 2.86% | 0.10% | 0.64% | 0.000% | 0.073% |
| Ascomycota | Eurotiomycetes | Eurotiales | Trichocomaceae | OTU 46 | 29 | 2.79% | 0.09% | 0.35% | 0.000% | 0.195% |
| Basidiomycota | Tremellomycetes | Cystofilobasidiales | Mrakiaceae | Udeniomyces | 19 | 2.71% | 0.09% | 0.45% | 0.000% | 0.295% |
| Pseudofungi | Oomycetes | Peronosporales | X | OTU 205 | 12 | 2.59% | 0.09% | 0.73% | 0.000% | 0.009% |
| Basidiomycota | Exobasidiomycetes | X | X | OTU 95 | 12 | 2.26% | 0.08% | 1.67% | 0.000% | 0.014% |
| Pseudofungi | Oomycetes | Peronosporales | Peronosporaceae IS | Phytophthora_C1c | 6 | 2.25% | 0.08% | 1.08% | 0.000% | 0.000% |
| Basidiomycota | Agaricomycetes | X | X | OTU 120 | 20 | 2.19% | 0.07% | 0.68% | 0.000% | 0.056% |
| Basidiomycota | Exobasidiomycetes | Exobasidiales | Exobasidiaceae | OTU 158 | 9 | 2.19% | 0.07% | 1.99% | 0.000% | 0.000% |
| Ascomycota | Saccharomycetes | Saccharomycetales | Saccharomycetaceae | Debaryomyces_marama | 30 | 2.18% | 0.07% | 0.40% | 0.005% | 0.075% |
| Pseudofungi | Oomycetes | Peronosporales | Peronosporaceae | Hyaloperonospora | 22 | 2.18% | 0.07% | 0.39% | 0.000% | 0.000% |
| Basidiomycota | Agaricostilbomycetes | Agaricostilbales | Agaricostilbaceae | Agaricostilbum | 13 | 2.17% | 0.07% | 0.87% | 0.000% | 0.007% |
| Ascomycota | Dothideomycetes | Pleosporales | X | OTU 3500 | 20 | 2.15% | 0.07% | 0.17% | 0.000% | 0.000% |
| Ascomycota | Dothideomycetes | Capnodiales | Capnodiales incertae sedi | Capnodiales incertae sedis | 13 | 2.05% | 0.07% | 0.59% | 0.000% | 0.006% |
| Ascomycota | Saccharomycetes | Saccharomycetales | Saccharomycetaceae | Lodderomyces_elongisporus | 29 | 2.05% | 0.07% | 0.25% | 0.000% | 0.054% |
| Ascomycota | Pezizomycetes | Pezizales | Ascodesmidaceae | Ascodesmis | 21 | 1.98% | 0.07% | 0.50% | 0.000% | 0.004% |
| Ascomycota | Dothideomycetes | Pleosporales | Phaeosphaeriaceae | Phaeodothis | 18 | 1.97% | 0.07% | 0.50% | 0.000% | 0.035% |
| Ascomycota | Dothideomycetes | X | X | OTU 2396 | 23 | 1.96% | 0.07% | 0.15% | 0.000% | 0.015% |
| Ascomycota | Saccharomycetes | Saccharomycetales | Saccharomycetaceae | Candida_catenulata_NRRL_Y-15( | 21 | 1.96% | 0.07% | 0.35% | 0.000% | 0.016% |
| Basidiomycota | Tremellomycetes | Tremellales | Bulleribasidiaceae | Dioszegia | 18 | 1.93% | 0.06% | 0.45% | 0.000% | 0.344% |
| Ascomycota | Dothideomycetes | Pleosporales | X | OTU 217 | 20 | 1.84% | 0.06% | 0.35% | 0.000% | 0.000% |
| Ascomycota | Eurotiomycetes | Eurotiales | Trichocomaceae | OTU 246 | 14 | 1.78% | 0.06% | 0.37% | 0.000% | 0.000% |
| Ascomycota | Dothideomycetes | Pleosporales | X | OTU 190 | 25 | 1.75% | 0.06% | 0.16% | 0.000% | 0.000% |
| Ascomycota | Leotiomycetes | Helotiales | Helotiaceae | Tricladium | 27 | 1.69% | 0.06% | 0.39% | 0.000% | 0.035% |
| Basidiomycota | Agaricomycetes | Agaricales | Entolomataceae | OTU 226 | 20 | 1.68% | 0.06% | 1.01% | 0.000% | 0.015% |
| Ascomycota | Saccharomycetes | Saccharomycetales | Saccharomycetaceae | Hyphopichia_burtonii_CBS_235 | 16 | 1.68% | 0.06% | 0.39% | 0.000% | 0.199% |
| Ascomycota | Saccharomycetes | Saccharomycetales | Saccharomycetaceae | Pichia_fermentans_(T) | 14 | 1.68% | 0.06% | 0.50% | 0.000% | 0.224% |
| Basidiomycota | Ustilaginomycetes | Ustilaginales | Ustilaginaceae | Pseudozyma | 18 | 1.66% | 0.06% | 0.27% | 0.000% | 0.000% |
| Fungi incertae sedi: | Mucoromycotina | Mucorales | Lichtheimiaceae | Lichtheimia | 21 | 1.65% | 0.05% | 1.24% | 0.000% | 0.014% |
| Ascomycota | Leotiomycetes | Helotiales | Dermateaceae | Mollisia | 15 | 1.59% | 0.05% | 0.60% | 0.000% | 0.041% |
| Basidiomycota | Cystobasidiomycetes | Erythrobasidiales | Erythrobasidiaceae | OTU 261 | 13 | 1.59% | 0.05% | 0.51% | 0.000% | 0.040% |
| Ascomycota | Dothideomycetes | Capnodiales | Mycosphaerellaceae | OTU 184 | 27 | 1.55% | 0.05% | 0.20% | 0.000% | 0.005% |
| Basidiomycota | Agaricomycetes | Polyporales | Lachnocladiaceae | Asterostroma | 8 | 1.54% | 0.05% | 0.60% | 0.000% | 0.000% |
| Ascomycota | Laboulbeniomycetes | Pyxidiophorales | Pyxidiophoraceae | Pyxidiophora | 14 | 1.54% | 0.05% | 0.56% | 0.000% | 0.036% |
| Basidiomycota | Exobasidiomycetes | Microstromatales | X | OTU 169 | 11 | 1.52% | 0.05% | 0.43% | 0.000% | 0.000% |
| Ascomycota | Dothideomycetes | Pleosporales | Didymellaceae | OTU 1903 | 24 | 1.52% | 0.05% | 0.11% | 0.000% | 0.011% |
| Basidiomycota | Exobasidiomycetes | Exobasidiales | Cryptobasidiaceae | Clinoconidium | 3 | 1.51% | 0.05% | 1.27% | 0.000% | 0.000% |
| X | X | X | X | OTU 248 | 4 | 1.49% | 0.05% | 0.52% | 0.000% | 0.000% |
| Ascomycota | Dothideomycetes | Pleosporales | Pleosporaceae | OTU 3417 | 11 | 1.48% | 0.05% | 0.38% | 0.000% | 0.012% |
| Ascomycota | Saccharomycetes | Saccharomycetales | Saccharomycetaceae | Issatchenkia_orientalis_ATCC_62 | 30 | 1.48% | 0.05% | 0.13% | 0.010% | 0.060% |
| Ascomycota | Dothideomycetes | X | X | OTU 231 | 17 | 1.47% | 0.05% | 0.25% | 0.000% | 0.000% |
| Ascomycota | Dothideomycetes | Pleosporales | Sporormiaceae | Preussia | 14 | 1.46% | 0.05% | 0.91% | 0.000% | 0.000% |
| Ascomycota | Dothideomycetes | Capnodiales | Dissoconiaceae | Dissoconium | 25 | 1.46% | 0.05% | 0.27% | 0.000% | 0.036% |
| Ascomycota | Dothideomycetes | Capnodiales | Capnodiales incertae sedi | Verrucocladosporium | 10 | 1.46% | 0.05% | 0.76% | 0.000% | 0.000% |
| Pseudofungi | Oomycetes | Albuginales | Albuginaceae | Albugo | 19 | 1.39% | 0.05% | 0.26% | 0.000% | 0.000% |
| Ascomycota | Saccharomycetes | Saccharomycetales | Saccharomycetaceae | Meyerozyma_guilliermondii | 15 | 1.36% | 0.05% | 0.40% | 0.000% | 0.064% |
| Ascomycota | Dothideomycetes | Pleosporales | Pleosporaceae | OTU 2324 | 22 | 1.36% | 0.05% | 0.10% | 0.000% | 0.006% |
| Ascomycota | Leotiomycetes | Helotiales | Sclerotiniaceae | Sclerotinia | 21 | 1.36% | 0.05% | 0.35% | 0.000% | 0.000% |
| Ascomycota | Dothideomycetes | Pleosporales | X | OTU 40 | 10 | 1.32% | 0.04% | 0.44% | 0.000% | 0.166% |
| Pseudofungi | Oomycetes | Peronosporales | Peronosporaceae | OTU 904 | 3 | 1.32% | 0.04% | 0.62% | 0.000% | 0.000% |
| Ascomycota | Eurotiomycetes | Eurotiales | Trichocomaceae | OTU 302 | 17 | 1.30% | 0.04% | 0.36% | 0.000% | 0.028% |
| Ascomycota | Leotiomycetes | Helotiales | X | OTU 121 | 11 | 1.28% | 0.04% | 0.35% | 0.000% | 0.057% |
| Ascomycota | Dothideomycetes | Pleosporales | Pleosporaceae | OTU 3654 | 21 | 1.28% | 0.04% | 0.11% | 0.000% | 0.006% |
| Ascomycota | Dothideomycetes | Pleosporales | X | OTU 2725 | 18 | 1.25% | 0.04% | 0.11% | 0.000% | 0.000% |
| Ascomycota | X | X | X | OTU 451 | 7 | 1.22% | 0.04% | 0.93% | 0.000% | 0.006% |
| Basidiomycota | Ustilaginomycetes | Ustilaginales | Ustilaginaceae | OTU 222 | 9 | 1.19% | 0.04% | 0.29% | 0.000% | 0.145% |
| Ascomycota | Dothideomycetes | Pleosporales | Pleosporaceae | OTU 2651 | 19 | 1.17% | 0.04% | 0.09% | 0.000% | 0.000% |
| Ascomycota | Dothideomycetes | Pleosporales | Pleosporales incertae sed | Camarosporium | 27 | 1.15% | 0.04% | 0.19% | 0.000% | 0.011% |
| Basidiomycota | Cystobasidiomycetes | Cystobasidiales | Cystobasidiaceae | Rhodotorula 1 -Cystobasidiales | 12 | 1.15% | 0.04% | 0.32% | 0.000% | 0.027% |
| Basidiomycota | Agaricomycetes | Agaricales | X | OTU 3288 | 13 | 1.15% | 0.04% | 0.50% | 0.000% | 0.000% |
| Ascomycota | Dothideomycetes | Pleosporales | X | OTU 1952 | 11 | 1.13% | 0.04% | 0.38% | 0.000% | 0.035% |
| Ascomycota | Dothideomycetes | Capnodiales | X | OTU 213 | 13 | 1.11% | 0.04% | 0.34% | 0.000% | 0.000% |
| Pseudofungi | Oomycetes | Peronosporales | Peronosporaceae | Bremia | 15 | 1.11% | 0.04% | 0.41% | 0.000% | 0.000% |
| Basidiomycota | Cystobasidiomycetes | Cystobasidiales | Cystobasidiaceae | Cystobasidium | 27 | 1.09% | 0.04% | 0.14% | 0.000% | 0.088% |
| Basidiomycota | Cystobasidiomycetes | Erythrobasidiales | Erythrobasidiaceae | OTU 301 | 20 | 1.07% | 0.04% | 0.15% | 0.000% | 0.000% |
| Ascomycota | Leotiomycetes | Thelebolales | Thelebolaceae | Thelebolus | 14 | 1.07% | 0.04% | 0.19% | 0.000% | 0.034% |
| Ascomycota | Dothideomycetes | Pleosporales | Pleosporaceae | OTU 5424 | 18 | 1.03% | 0.03% | 0.09% | 0.000% | 0.000% |
| Entomophthoromy | Entomophthoromycete | Entomophthorales | Ancylistaceae | Conidiobolus | 7 | 1.02% | 0.03% | 0.36% | 0.000% | 0.000% |
| Fungi incertae sedi: | Mortierellomycotina | Mortierellales | Mortierellaceae | Mortierella | 22 | 1.02% | 0.03% | 0.18% | 0.000% | 0.000% |
| Ascomycota | Dothideomycetes | Botryosphaeriales | Botryosphaeriaceae | Lasiodiplodia | 21 | 1.01% | 0.03% | 0.10% | 0.000% | 0.000% |
| Basidiomycota | Exobasidiomycetes | Tilletiales | Tilletiaceae | Tilletia | 25 | 1.00% | 0.03% | 0.12% | 0.000% | 0.103% |
| Fungi incertae sedi: | Mucoromycotina | Mucorales | Mucoraceae | OTU 281 | 7 | 1.00% | 0.03% | 0.92% | 0.000% | 0.006% |
| Ascomycota | Dothideomycetes | Pleosporales | X | OTU 4751 | 19 | 1.00% | 0.03% | 0.08% | 0.000% | 0.004% |
| Ascomycota | Eurotiomycetes | Chaetothyriales | Herpotrichiellaceae | OTU 610 | 6 | 1.00% | 0.03% | 0.84% | 0.000% | 0.000% |
| Basidiomycota | Cystobasidiomycetes | Erythrobasidiales | Erythrobasidiaceae | Sporobolomyces | 29 | 0.99% | 0.03% | 0.09% | 0.000% | 0.038% |
| Basidiomycota | Agaricomycetes | Polyporales | Coriolaceae | Bjerkandera | 11 | 0.98% | 0.03% | 0.35% | 0.000% | 0.004% |
| Ascomycota | X | X | X | OTU 537 | 7 | 0.96% | 0.03% | 0.48% | 0.000% | 0.004% |
| Ascomycota | Dothideomycetes | Pleosporales | X | OTU 1197 | 10 | 0.95% | 0.03% | 0.35% | 0.000% | 0.007% |
| Ascomycota | Dothideomycetes | Pleosporales | Pleosporaceae | OTU 5572 | 11 | 0.92% | 0.03% | 0.26% | 0.000% | 0.004% |
| Ascomycota | X | X | X | OTU 241 | 20 | 0.90% | 0.03% | 0.19% | 0.000% | 0.000% |
| Basidiomycota | Agaricomycetes | Agaricales | Clavariaceae | Clavaria_CVZO | 13 | 0.89% | 0.03% | 0.27% | 0.000% | 0.035% |
| Ascomycota | Dothideomycetes | Pleosporales | Pleosporaceae | Pyrenochaeta | 28 | 0.88% | 0.03% | 0.13% | 0.000% | 0.015% |
| Ascomycota | Eurotiomycetes | Eurotiales | Trichocomaceae | OTU 371 | 26 | 0.88% | 0.03% | 0.15% | 0.000% | 0.012% |
| Basidiomycota | Agaricomycetes | Hymenochaetales | X | OTU 105 | 20 | 0.88% | 0.03% | 0.15% | 0.000% | 0.012% |
| Ascomycota | Saccharomycetes | Saccharomycetales | Saccharomycetaceae | Saccharomyces_kudriavzevii | 28 | 0.87% | 0.03% | 0.16% | 0.000% | 0.022% |
| Ascomycota | Leotiomycetes | Erysiphales | Erysiphaceae | OTU 251 | 9 | 0.86% | 0.03% | 0.23% | 0.000% | 0.000% |
| Basidiomycota | Exobasidiomycetes | Microstromatales | Quambalariaceae | Quambalaria | 20 | 0.84% | 0.03% | 0.14% | 0.000% | 0.000% |
| Basidiomycota | Microbotryomycetes | Microbotryomycetes | Microbotryomycetes inc | Kriegeria | 15 | 0.82% | 0.03% | 0.21% | 0.000% | 0.039% |
| Ascomycota | Saccharomycetes | Saccharomycetales | Saccharomycetaceae | Candida_intermedia_CBS_7153_ | 19 | 0.82% | 0.03% | 0.28% | 0.000% | 0.079% |
| Ascomycota | Leotiomycetes | Helotiales | Sclerotiniaceae | OTU 268 | 4 | 0.81% | 0.03% | 0.26% | 0.000% | 0.000% |
| Ascomycota | X | X | X | OTU 1279 | 1 | 0.78% | 0.03% | 0.78% | 0.000% | 0.000% |
| Basidiomycota | Agaricomycetes | Cantharellales | Ceratobasidiaceae | Thanatephorus | 19 | 0.77% | 0.03% | 0.35% | 0.000% | 0.000% |
| Ascomycota | Eurotiomycetes | Eurotiales | Trichocomaceae | OTU 763 | 23 | 0.77% | 0.03% | 0.10% | 0.000% | 0.012% |
| Ascomycota | Eurotiomycetes | Chaetothyriales | Herpotrichiellaceae | Sorocybe | 15 | 0.77% | 0.03% | 0.18% | 0.000% | 0.000% |
| Basidiomycota | Cystobasidiomycetes | Erythrobasidiales | Erythrobasidiaceae | Rhodotorula 2 -Erythrobasidiales | 13 | 0.77% | 0.03% | 0.17% | 0.000% | 0.074% |
| Ascomycota | Dothideomycetes | Pleosporales | Lophiostomataceae | Lophiostoma | 8 | 0.73% | 0.02% | 0.55% | 0.000% | 0.000% |
| Ascomycota | Saccharomycetes | Saccharomycetales | Saccharomycetaceae | OTU 253 | 6 | 0.73% | 0.02% | 0.34% | 0.000% | 0.342% |
| Ascomycota | Dothideomycetes | Capnodiales | Mycosphaerellaceae | OTU 447 | 6 | 0.73% | 0.02% | 0.53% | 0.000% | 0.005% |
| Basidiomycota | Agaricomycetes | Agaricales | X | OTU 6574 | 9 | 0.72% | 0.02% | 0.42% | 0.000% | 0.000% |
| Basidiomycota | Microbotryomycetes | X | X | OTU 1557 | 9 | 0.71% | 0.02% | 0.25% | 0.000% | 0.175% |
| Ascomycota | Dothideomycetes | X | X | OTU 2825 | 19 | 0.70% | 0.02% | 0.06% | 0.000% | 0.005% |
| Ascomycota | Dothideomycetes | Capnodiales | Capnodiales incertae sedi | Toxicocladosporium | 23 | 0.70% | 0.02% | 0.15% | 0.000% | 0.147% |
| Ascomycota | Dothideomycetes | Capnodiales | Capnodiales incertae sedi | OTU 509 | 10 | 0.69% | 0.02% | 0.27% | 0.000% | 0.121% |
| Ascomycota | Dothideomycetes | X | X | OTU 3834 | 18 | 0.68% | 0.02% | 0.08% | 0.000% | 0.000% |
| Ascomycota | Dothideomycetes | Pleosporales | Didymellaceae | OTU 5573 | 21 | 0.67% | 0.02% | 0.06% | 0.000% | 0.000% |
| Ascomycota | Dothideomycetes | Pleosporales | Didymellaceae | OTU 3531 | 29 | 0.67% | 0.02% | 0.08% | 0.000% | 0.029% |
| Ascomycota | Dothideomycetes | Pleosporales | X | OTU 4476 | 17 | 0.67% | 0.02% | 0.10% | 0.000% | 0.000% |
| X | X | X | X | OTU 74 | 3 | 0.66% | 0.02% | 0.45% | 0.000% | 0.016% |
| Ascomycota | Eurotiomycetes | Chaetothyriales | Herpotrichiellaceae | OTU 309 | 22 | 0.66% | 0.02% | 0.21% | 0.000% | 0.019% |
| Ascomycota | Dothideomycetes | Pleosporales | X | OTU 346 | 22 | 0.66% | 0.02% | 0.09% | 0.000% | 0.000% |
| Ascomycota | Ascomycota incertae se | Ascomycota incertae | Ascomycota incertae sedi | Radulidium | 5 | 0.66% | 0.02% | 0.63% | 0.000% | 0.004% |
| Ascomycota | Dothideomycetes | Capnodiales | Capnodiaceae | Microxyphium | 2 | 0.66% | 0.02% | 0.65% | 0.000% | 0.000% |
| Basidiomycota | Agaricomycetes | Agaricales | X | OTU 5955 | 11 | 0.65% | 0.02% | 0.37% | 0.000% | 0.000% |
| Ascomycota | Dothideomycetes | Pleosporales | Didymellaceae | OTU 4539 | 24 | 0.65% | 0.02% | 0.05% | 0.000% | 0.000% |
| Ascomycota | Dothideomycetes | Pleosporales | Pleosporaceae | OTU 3146 | 18 | 0.64% | 0.02% | 0.05% | 0.000% | 0.000% |
| Ascomycota | X | X | X | OTU 79 | 16 | 0.63% | 0.02% | 0.12% | 0.000% | 0.043% |
| Ascomycota | Dothideomycetes | Pleosporales | X | OTU 437 | 7 | 0.62% | 0.02% | 0.23% | 0.000% | 0.006% |
| Basidiomycota | Agaricomycetes | Agaricales | Hygrophoraceae | Cuphophyllus1_pratensis | 18 | 0.62% | 0.02% | 0.26% | 0.000% | 0.004% |
| Ascomycota | X | X | X | OTU 677 | 4 | 0.60% | 0.02% | 0.25% | 0.000% | 0.000% |
| Ascomycota | Eurotiomycetes | Eurotiales | Trichocomaceae | OTU 504 | 12 | 0.60% | 0.02% | 0.15% | 0.000% | 0.145% |
| Basidiomycota | Tremellomycetes | Holtermanniales | HoltermannialesFAM | Holtermanniella | 10 | 0.60% | 0.02% | 0.22% | 0.000% | 0.075% |
| Basidiomycota | Ustilaginomycetes | Urocystales | Urocystaceae | Urocystis | 13 | 0.59% | 0.02% | 0.18% | 0.000% | 0.000% |
| Basidiomycota | Exobasidiomycetes | Microstromatales | Microstromataceae | Microstroma | 22 | 0.59% | 0.02% | 0.15% | 0.000% | 0.013% |
| Ascomycota | Eurotiomycetes | Eurotiales | Trichocomaceae | OTU 2048 | 4 | 0.59% | 0.02% | 0.35% | 0.000% | 0.000% |
| Ascomycota | Pezizomycetes | Pezizales | Ascobolaceae | Ascobolus | 20 | 0.59% | 0.02% | 0.27% | 0.000% | 0.004% |
| Ascomycota | Dothideomycetes | Pleosporales | X | OTU 2945 | 12 | 0.58% | 0.02% | 0.11% | 0.000% | 0.047% |
| Basidiomycota | Agaricomycetes | X | X | OTU 436 | 9 | 0.58% | 0.02% | 0.23% | 0.000% | 0.231% |
| Ascomycota | Dothideomycetes | Pleosporales | X | OTU 5417 | 18 | 0.58% | 0.02% | 0.05% | 0.000% | 0.000% |
| Ascomycota | Saccharomycetes | Saccharomycetales | Saccharomycetaceae | OTU 1228 | 2 | 0.58% | 0.02% | 0.33% | 0.000% | 0.000% |
| Basidiomycota | Agaricomycetes | Agaricales | Agaricales incertae sedis | Agaricales | 3 | 0.58% | 0.02% | 0.55% | 0.000% | 0.014% |
| X | X | X | X | OTU 653 | 1 | 0.57% | 0.02% | 0.57% | 0.000% | 0.000% |
| X | X | X | X | OTU 466 | 8 | 0.56% | 0.02% | 0.32% | 0.000% | 0.046% |
| Ascomycota | Saccharomycetes | Saccharomycetales | Saccharomycetaceae | Torulaspora_pretoriensis | 23 | 0.55% | 0.02% | 0.08% | 0.000% | 0.015% |
| Ascomycota | Dothideomycetes | Pleosporales | Pleosporaceae | OTU 5843 | 19 | 0.55% | 0.02% | 0.04% | 0.000% | 0.000% |
| Ascomycota | Dothideomycetes | Pleosporales | X | OTU 3501 | 18 | 0.55% | 0.02% | 0.07% | 0.000% | 0.000% |
| Ascomycota | Dothideomycetes | Pleosporales | X | OTU 378 | 21 | 0.54% | 0.02% | 0.11% | 0.000% | 0.000% |
| Ascomycota | Saccharomycetes | Saccharomycetales | Saccharomycetaceae | Pichia_manshurica | 11 | 0.54% | 0.02% | 0.36% | 0.000% | 0.005% |
| Ascomycota | Eurotiomycetes | Chaetothyriales | Herpotrichiellaceae | [illegible] | [illegible] | [illegible] | [illegible] | [illegible] | [illegible] | [illegible] |
